# Post-trauma behavioral phenotype predicts vulnerability to fear relapse after extinction

**DOI:** 10.1101/2021.09.25.461769

**Authors:** Fanny Demars, Ralitsa Todorova, Gabriel Makdah, Antonin Forestier, Marie-Odile Krebs, Bill P. Godsil, Thérèse M. Jay, Sidney I. Wiener, Marco N. Pompili

## Abstract

Current treatments for trauma-related disorders remain ineffective for many patients. Here, we modeled interindividual differences in post-therapy fear relapse with a novel ethologically relevant trauma recovery paradigm. After traumatic fear conditioning, male rats underwent fear extinction while foraging in a large enriched arena, permitting the expression of a wide spectrum of behaviors, assessed by an automated pipeline. This multidimensional behavioral assessment revealed that post-conditioning fear response profiles clustered into two groups, respectively characterized by active vs. passive fear responses. After trauma, some animals expressed fear by freezing, while others darted, as if fleeing from danger. Remarkably, belonging to the darters or freezers group predicted differential levels of vulnerability to fear relapse after extinction. Moreover, genome-wide transcriptional profiling revealed that these groups differentially regulated specific sets of genes, some of which have previously been implicated in anxiety and trauma-related disorders. Our results suggest that post-trauma behavioral phenotypes and the associated epigenetic landscapes can serve as markers of fear relapse susceptibility, and thus may be instrumental for future development of more effective treatments for psychiatric patients.

## INTRODUCTION

Anxiety and trauma-related disorders constitute major public health challenges. However, available treatments remain only partially effective (Hoskins et al. 2015; Cusack et al. 2016). Fear extinction deficiency is a prominent feature of these diseases (Mary et al. 2020), and many behavioral treatments, such as exposure-based therapies (EBT), rely on extinction training (Milad et al. 2009; Powers et al. 2010). However, in many patients, even successful EBT in the clinic is followed by a relapse of symptoms in the course of the patients’ daily lives. The pathophysiological mechanisms underlying such interindividual variations in vulnerability to relapse remain unknown (Etkin et al. 2019; Zhutovsky et al. 2019; Korgaonkar et al. 2020), and a crucial endeavor in fear extinction research is to identify clear biomarkers of such differences.

Rodent experimental models of fear behavior, and their neurobiological substrates, translate well to humans (Mobbs et al. 2007). Thus, fear conditioning has been used for decades to study the biological correlates of aversive learning and memory in human and rodents. In particular, context-dependent fear renewal paradigms in rodents are employed to model fear relapse after therapy (e.g. Marek et al. 2018). In order to model the propensity for pathological responses in humans, it is necessary to characterize animals’ inter-individual differences (Holmes and Singewald 2013; Headley et al. 2019). Moreover, some biomarkers may only appear when data is partitioned according to the particular response phenotypes of individuals (Cohen and Zohar 2004; Peters et al. 2010; Dopfel et al. 2019). Indeed, interindividual differences in conditioned fear responses, extinction, and relapse behaviors have been observed in animals (Milad and Quirk 2002; Bush et al. 2007; Duvarci et al. 2009; Peters et al. 2010; Galatzer-Levy et al. 2013; Reznikov et al. 2015; Gruene et al. 2015). However, despite the evidence of significant between-subject behavioral variability during extinction learning, reliable predictive behavioral markers of vulnerable vs. resilient fear relapse phenotypes are still lacking.

While traditionally, freezing behavior is employed as the sole indicator of fear measured in conditioning and extinction protocols, recent work has also investigated active fear responses after fear conditioning in the form of flight-like behavior, revealing differences between individuals in their propensity for passive vs. active fear responses (Gruene et al. 2015; Fadok et al. 2017; Totty et al. 2021). Such interindividual differences may be useful indicators of the vulnerability to fear relapse after extinction. Nevertheless, a shortcoming of assessing animal behavior in standard conditioning chambers (Skinner boxes) is the restricted behavioral repertoire that can be expressed there, potentially reducing detectible differences between individuals. On the other hand, more ecologically relevant settings would allow expression of a broader repertoire of animals’ natural behavioral patterns (Krakauer et al. 2017), therefore better modelling and contrasting healthy and pathological behavioral profiles. Recent years have seen a surge in novel naturalistic approaches to study fear behavior (Mobbs and Kim 2015; Paré and Quirk 2017; Kim and Jung 2018; Headley et al. 2019). But these advances have not yet revealed behavioral and biological markers of interindividual differences in the vulnerability to fear relapse after extinction.

Here, we hypothesized that soon following trauma, diverse behavioral phenotypes may be unveiled in an enriched naturalistic setting permitting the animals to express a wide spectrum of behaviors. We also hypothesized that these early behavioral markers may predict individual vulnerability to fear relapse. To test this, we developed a multidimensional automated behavioral assessment pipeline, and this revealed two distinct fear response profiles during early extinction training. Remarkably, these post-conditioning phenotypes predicted animals’ vulnerability to fear relapse intensity. Moreover, a genome-wide transcriptional profiling of the ventromedial prefrontal cortex (vmPFC), critically implicated in extinction, renewal, and PTSD (Milad and Quirk 2002; Kalisch et al. 2006; Milad et al. 2007; Peters et al. 2010; Sierra-Mercado et al. 2011; Stafford et al. 2012; Knapska et al. 2012; Garfinkel et al. 2014; Do-Monte et al. 2015; Marek et al. 2018), showed that, beyond behavior, these phenotypes were also characterized by differential biological substrates. Indeed, the two groups of animals also displayed divergent gene expression profiles, including genes previously implicated in anxiety and trauma-related disorders, pointing toward potential molecular substrates of differential fear renewal vulnerability.

## RESULTS

### Complex behavioral patterns in an ethologically relevant fear extinction paradigm

We designed a new, ethologically rich model of trauma recovery where fear extinction takes place in a large open field arena. Male rats were habituated to forage for food pellets in the arena enriched with large objects, thus permitting the animals to display a wide spectrum of behaviors. Then rats underwent a typical fear conditioning protocol in a standard apparatus, followed by fear extinction training in the large arena (**Fig. 1a**). We used head movements, orientation and position measures, and machine learning, to automatically classify animal behavior (**Fig. 1b**) in six classes: freezing, darting, grooming, object exploration, rearing, and all remaining foraging and exploratory behaviors, referred to collectively as foraging/exploration. Darting is a behavior where animals run at high speeds in long straight trajectories, as already observed in extended environments (Reinhold et al. 2019) (**Fig. 1c-d**). This contrasted with the locomotor patterns associated with foraging, where the animals moved about at low to medium speeds in various directions (**Supp. Video 1**). Since average walking and running speed varied across animals, a specific criterion was identified for each individual with unsupervised clustering of its speed distribution (see Methods).

**Figure 1.**
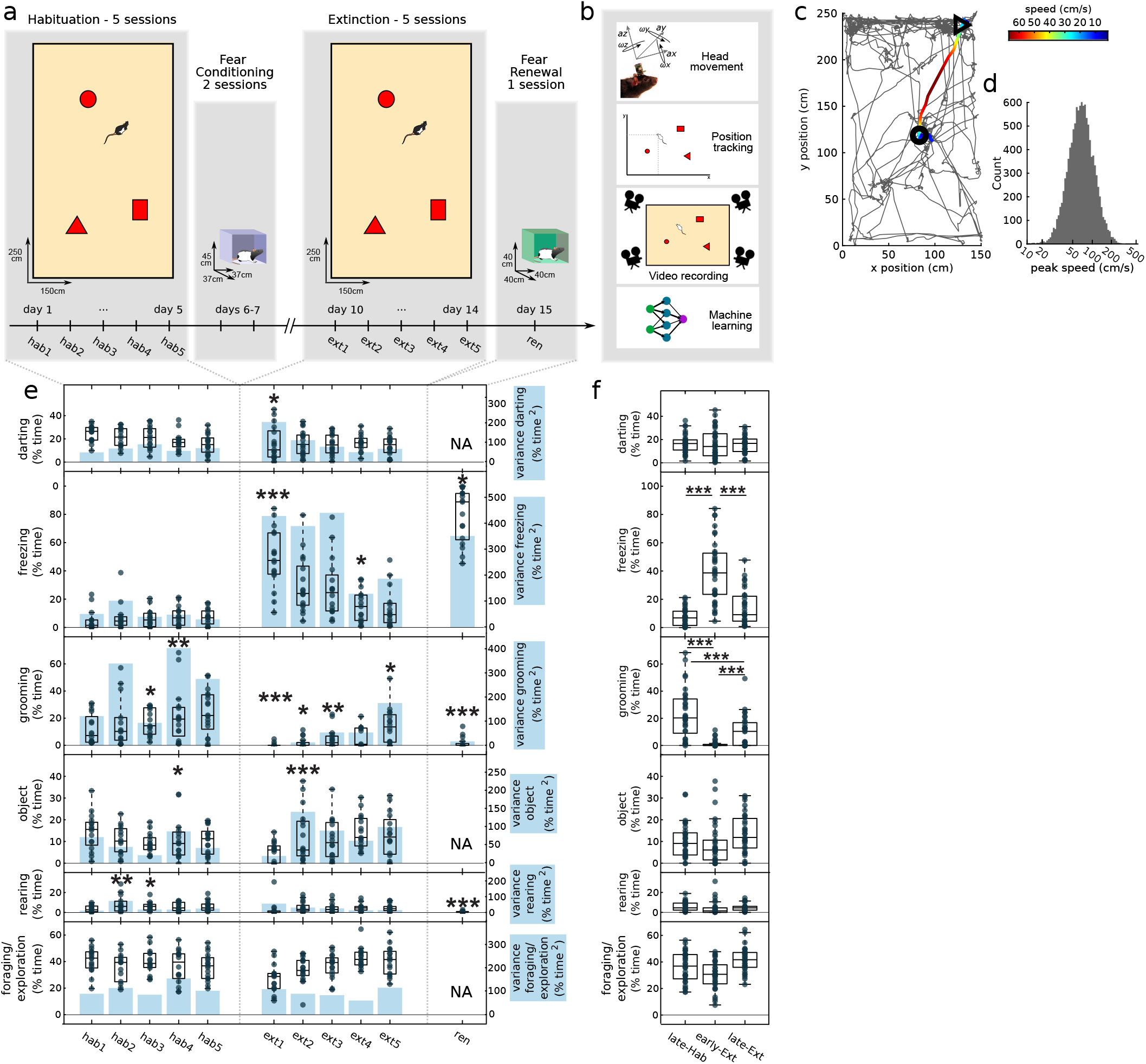
Multidimensional behavioral scoring in an ethologically-relevant paradigm. (**a**) Behavioral protocol. Eighteen rats were habituated to forage in a large open field with sheltering objects (red shapes) for 5 days (hab1 to hab5). Then they underwent an auditory fear conditioning protocol for two days in a standard conditioning cubicle. Five days of extinction (ext) training took place in the open field while the rats foraged, followed by the fear renewal test (ren). (**b**) Automated multidimensional behavioral assessment with head movement data (3D accelerations and rotations), position tracking, recorded videos, and machine learning. (**c**) Example of a darting trajectory. Overhead view of the open field environment with gray traces depicting the trajectories of the animal over 5 min. The darting trajectory is highlighted with the thick colored line. Colors indicate speed of the animal. Triangle: darting trajectory start; circle: darting trajectory end. (**d**) Distribution of peak speeds of all detected darting trajectories of all rats. (**e**) Time spent performing behaviors during CS periods and group variance. Dots represent individual rats’ averages over the three CS periods. In box plots, the central bar indicates the median, the bottom and top edges indicate the 25th and 75th percentiles; whiskers extend to the most extreme data points, excluding outliers (see Methods). The blue shading bars depict variance (scales at right). Stars mark the sessions where the variance was different from the previous session (F-test for equal variances). Note that object exploration, darting, and other exploratory/foraging behavior could not be expressed (NA) in the small conditioning chamber. (**f**) Comparison of behavioral expression during late habituation (late-Hab: hab4 and hab5), early extinction (early-Ext: ext1 and ext2), and late extinction (late-Ext: ext4 and ext5) sessions. Stars denote significant differences between these training stages (sign rank test). Only significant comparisons are shown. [*p<0.05, **p<0.01, ***p<0.001]

Following auditory fear conditioning, during fear extinction training in the open field, we observed the expected freezing responses to the conditioned stimulus (CS). Freezing to the CS was initially higher than during habituation, but, after three days of cued fear extinction training in the open field, it was indistinguishable from baseline (**Fig. 1e-f**). This demonstrated that cued conditioned fear was also recalled in the open field context, and that this fear successfully reduced with extinction training. Other behaviors were also affected after fear conditioning, such as grooming, which was reduced (**Fig. 1f**). Importantly, behavioral changes were not uniform among animals and, notably, interindividual variability increased in freezing and darting (**Fig. 1e**), suggesting that these behaviors may constitute useful targets for post-trauma behavioral profiling.

### Interindividual differences in fear responses during extinction training

Overall, during early extinction, CS presentations elicited highly variable behavioral responses among animals (**Fig. 2a**). We hypothesized that this variability corresponded to diverging behavioral phenotypes, which, in turn, would correspond to differing degrees of susceptibility to relapse after extinction. Principal component analysis (PCA) of the expression of the six classes of behavior after CS presentations aimed to characterize this variability of fear responses during early extinction training in an unbiased manner, accounting for all the behavioral variance in our data. The first principal component (PC1) explained most of the variance (**Fig. 2b**) and was characterized by opposing contributions of darting and freezing (**Fig. 2c**). By clustering the expression of the three significant PCs in the first extinction session (ext1), animals were divided into distinct groups (**Fig. 2d**, **Supp. Fig. 1a-b**), without any a priori assumption about which specific behavior would better define behavioral phenotypes. This revealed that the two groups differed mainly by the amount of freezing and darting in early extinction training (**Fig. 3a-b**, **Supp. Figs. 1c and 2**). Notably, these behavioral profiles emerged on the first extinction session after fear conditioning, and reconverged with extinction training (**Fig. 3g-j**), further suggesting that behavioral divergence originated from the traumatic experience. We refer to these two groups as *freezers* and *darters* according to their respective dominant CS-evoked responses in early extinction sessions.

**Figure 2.**
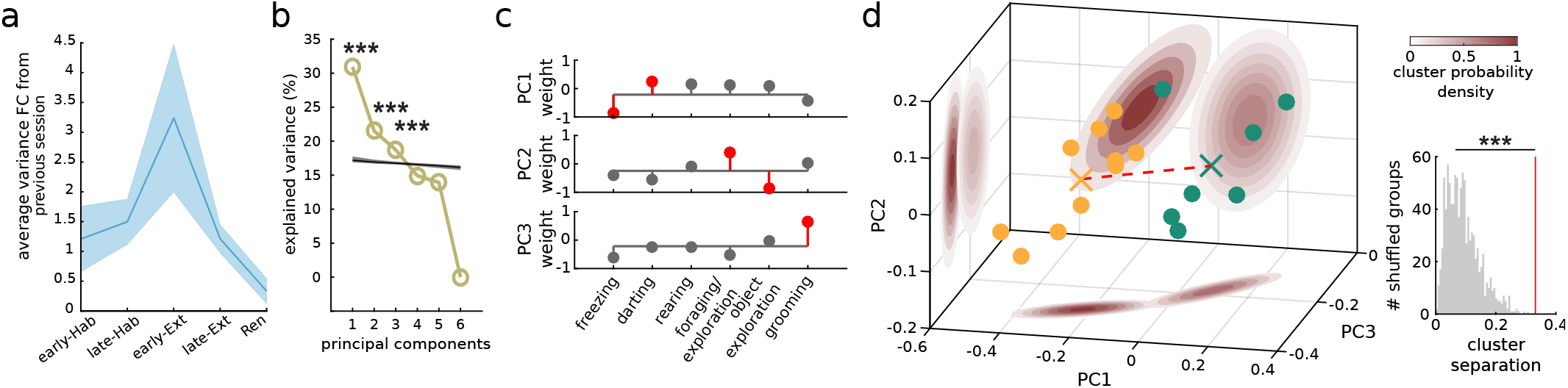
Analysis of the multidimensional behavioral data space to cluster distinct post-trauma fear response profiles. (**a**) Fold change (FC) in average variance of all behavior classes (pooled) compared to the previous session at different training stages (early-Hab and early-Ext includes only hab2 and ext2 respectively since the change compared to previous day cannot be computed for hab1 and ext1). Solid line: mean; shading: SEM. (**b**) Proportion of the variance explained by each PC. Black line: mean explained variance obtained for shuffled data (shading: SEM). Stars denote components whose explained variance is significantly different from chance (Monte Carlo bootstrap, *** p<0.001). (**c**) Respective contributions of each behavioral class to the first three PCA components. Red indicate significant weights (Otsu’s treshold, see Methods). (**d**) Expression of significant PCs during CS periods in ext1. Dots represent averages for the three CS for each rat. Two groups of individuals, yellow and green dots, are separated by k-means clustering. Xs: centers of mass of the two clusters; red dashed line: their Euclidean distance; red shaded areas: contour lines of the Gaussian model estimating densities; inset: distribution of the inter-cluster distances obtained for 1000 shuffles of group membership (red vertical line: actual distance between the two groups which is beyond the 99th percentile). [***p<0.001]

**Figure 3.**
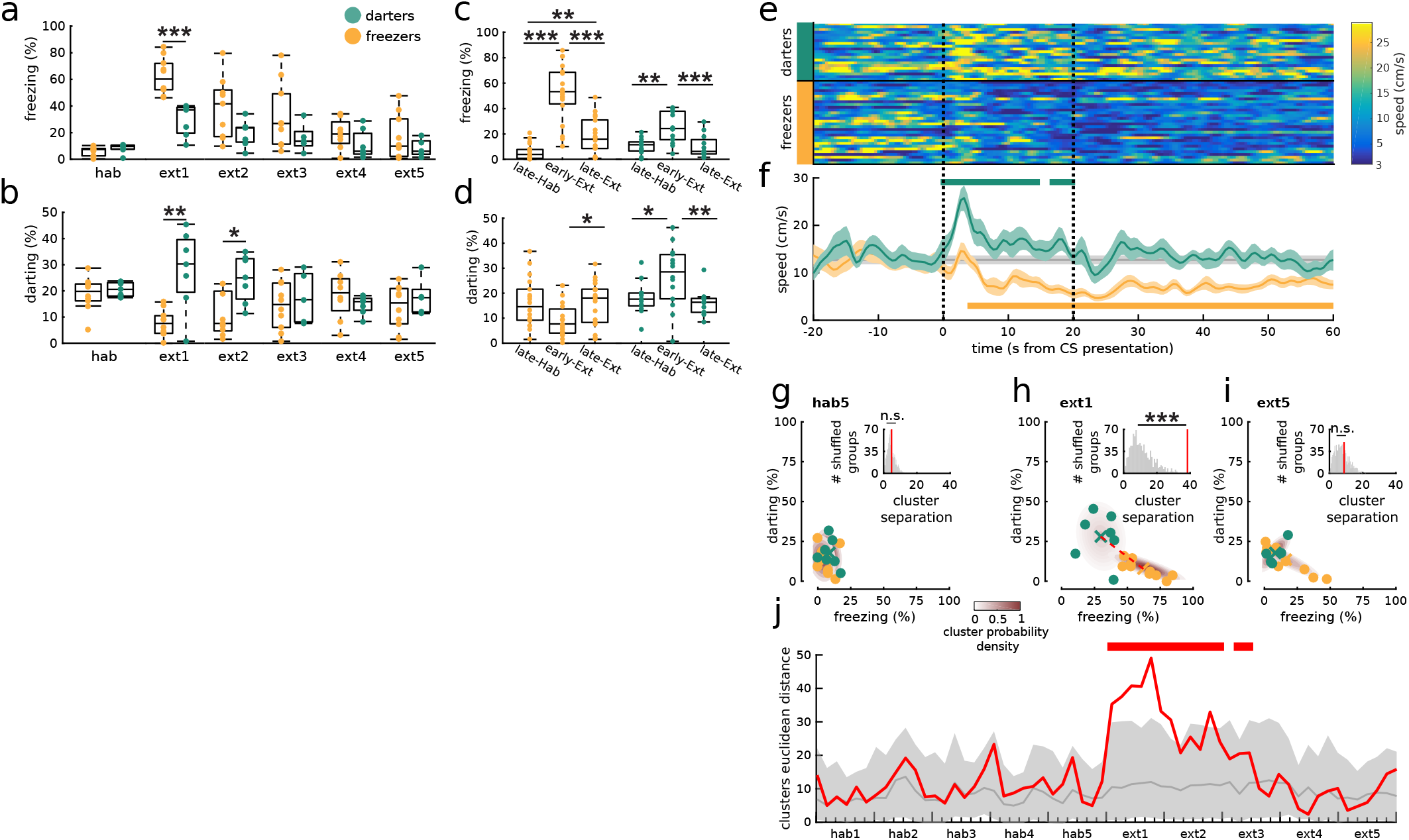
Distinct fear response profiles and their evolution over extinction training. Intergroup comparisons of freezing (**a**) and darting (**b**). Dots: individual rats’ averages over the three CS periods. Box plot format is the same as in Fig. 1c. (**c-d**) Intra-group comparisons of darting or freezing over extinction training (grouping as in Fig. 1d). (a,b,c,d) sign-rank tests. (**e-f**) Locomotion speed around all CS presentations on ext1 for the two groups. Black dashed vertical lines: CS onset and offset. (e) All individual trials. (f) Averages (solid lines) and SEM (shading) of values in (e) for darters and freezers. Gray bar: average baseline speeds (over the 20 s prior to CS onset) for all animals. Horizontal bars above and below denote periods when speeds were significantly different from baseline for darters and freezers, respectively (unpaired t-test). (**g-i**) After trauma (h), darters and freezers formed two distinct clusters in terms of their propensity to dart or freeze after CS presentation; no difference was observed before conditioning (g) or at the end of extinction training (i). Similar format to Fig. 2d. (**j**) Distance between darters and freezers in the darting vs freezing space (like panels g-i) over time (red curve) compared to shuffled groups (gray line: mean; shading: 95% confidence interval). Red bars above: periods when clusters separation was greater than chance (Monte Carlo bootstrap). [*p<0.05, **p<0.01, ***p<0.001]

One possible source of differences in behavioral responses to the CS during extinction could have been intergroup differences in fear conditioning or contextual fear generalization (from the conditioning box to the open field). However, no difference between groups was observed during fear acquisition (**Supp. Fig. 3a**). Moreover, freezing levels were low during the ext1 baseline period prior to CS presentation, suggesting weak contextual fear, and were not significantly different between groups (**Supp. Fig. 3b**). Another possible confound could be inter-group differences in within-day extinction learning. But, the difference of freezing levels between the last and the first CS presentations was not significantly different between the two groups (**Supp. Fig. 3c**). As a control, the clustering analysis was performed with data from the last day of extinction, but this did not lead to groups with significant differences in behavior evolution over extinction (**Supp. Fig. 1d-f**), indicating that interindividual differences in extinction behaviors were best captured by the phenotypes expressed in early extinction.

Like freezing, darting behavior increased after conditioning and decreased over extinction training in darters (**Fig. 3c-d**). Moreover, darting was evoked by CS presentations (**Fig. 3e-f**). These results are consistent with darting as an expression of conditioned defensive behavior specific for a subpopulation of the animals. Indeed, darting may be considered as an active behavioral pattern resembling flight and avoidance responses (Choi and Kim 2010; Gruene et al. 2015; Fadok et al. 2017; Kyriazi et al. 2018), since darting trajectories principally started and ended at sheltering locations in the arena (**Supp. Fig. 4a-b**). Moreover, there was no significant difference in distance from sheltering locations between darters and freezers at CS onset (**Supp. Fig. 4c**), ruling out the hypothesis that freezers darted less because they were already at sheltering locations at CS onset.

### Post-trauma behavioral phenotypes predict differential levels of vulnerability to fear renewal

We then quantified fear renewal to test the hypothesis that post-trauma behavioral profiles would predict contextdependent traumatic fear relapse. After 5 days of extinction training in the open field, the animals were placed in a standard conditioning chamber similar in size to those where initial conditioning took place, but different enough to be successfully discriminated, since it evoked low contextual freezing levels (**Fig. 4a**). As expected, CS presentation there elicited robust fear renewal in the form of high CS-triggered freezing relative to levels at the end of extinction training (**Fig. 1e**). Surprisingly, here the CS triggered significantly more freezing in darters than in freezers (**Fig. 4b**). This difference in fear renewal response strengths could not be accounted for by differences between groups in contextual fear levels or extinction rates during the renewal test (**Fig. 4a,c**).

**Figure 4.**
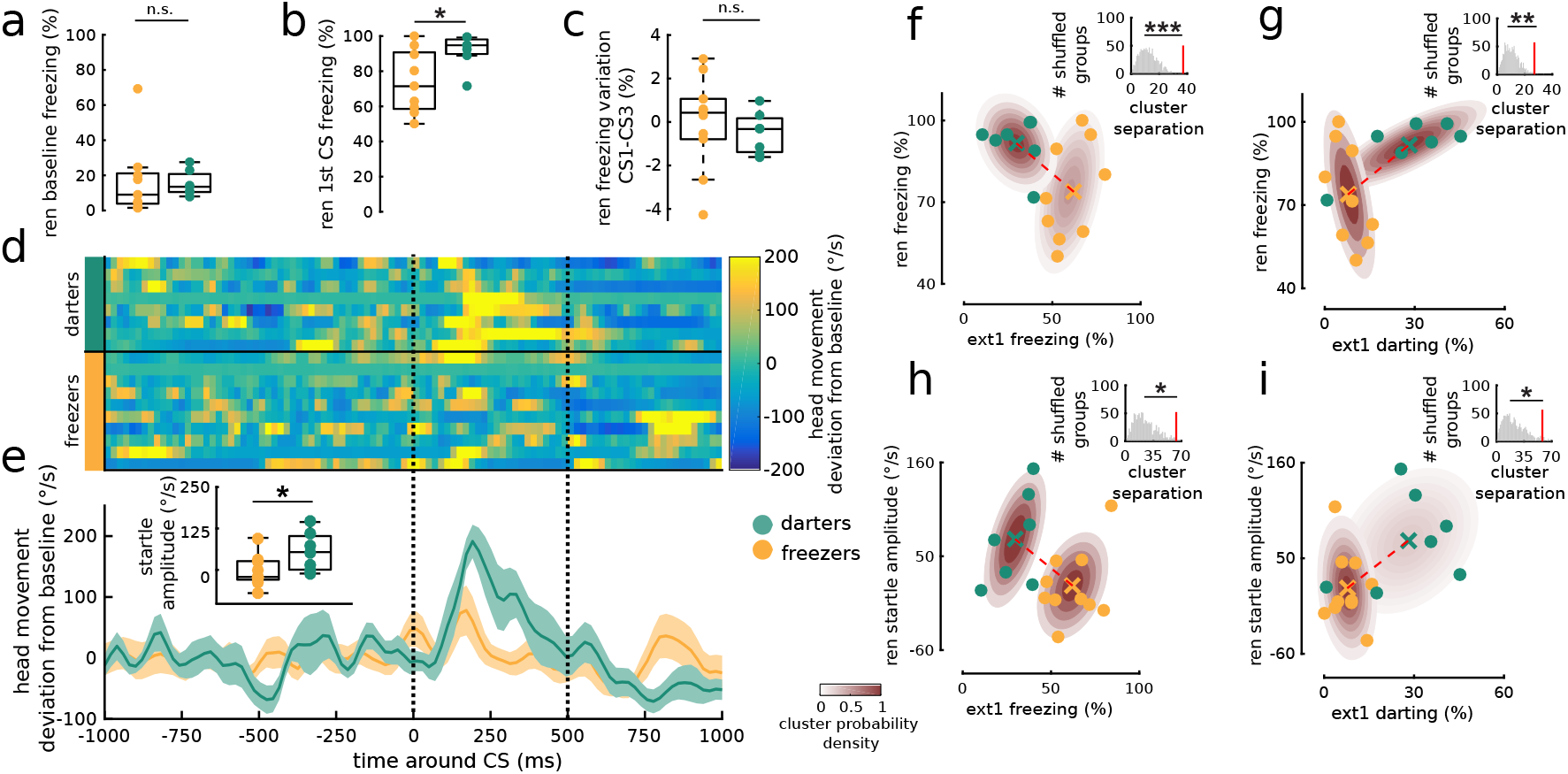
Differential vulnerability to fear renewal in darters vs. freezers. (**a**) Time spent freezing during the baseline period prior to the first CS presentation. (**b**) Freezing during the first CS period is significantly greater in darters. (**c**) Within-session learning rate expressed as the difference in freezing between the first and final CS periods (unpaired t-tests). (**d-e**) Orienting responses measured from head angular velocity. (d) Individual trials. (e) Average (solid lines) and SEM (shading) for darters and freezers. The inset quantifies the startle amplitude defined as the average head movement velocity during the first 500 ms after CS onset. Note that darters manifested a significantly stronger CS-evoked acoustic startle response (unpaired t-test). (**f-i**) Fear renewal strength, measured as freezing (f-g) or startle (h-i) to the first CS presentation, as a function of the amount of darting or freezing after trauma, during ext1. (insets) Darters and freezers formed two distinct clusters in terms of renewal freezing vs. ext1 freezing (f) or vs. ext1 darting (g) as well as of renewal startle vs. ext1 freezing (h) or darting (i). Same format and statistics as in Fig. 3g-i. [n.s.: not significant, *p<0.05, **p<0.01, ***p<0.001]

Complementing freezing measures, we quantified the magnitude of the orienting response the rats displayed upon CS onset (**Supp. Video 2**) by measuring head movement. We took this as a proxy of the magnitude of the acoustic startle response, which is known to be potentiated by fear (Davis et al. 1993), although our setup was not suitable to measure the starle reflex itself. Orienting magnitude too was greater in darters than freezers during the renewal test (**Fig. 4d-e**), corroborating the result for freezing. Overall, the tendency of an animal to dart or to freeze after CS presentation during the first extinction session in a large open field was respectively associated with higher or lower vulnerability to context-dependent fear renewal (**Fig. 4f-i**).

### Fear renewal vulnerability phenotypes are characterized by different gene expression profiles

Finally, we sought to characterize specific biological substrates distinguishing these two behavioral phenotypes. Epigenetic mechanisms have been associated with the recall and extinction of conditioned fear (Soliman et al. 2010; Lin et al. 2011; Stafford et al. 2012; Li et al. 2019), as well as with Post-Traumatic Stress Disorder (PTSD) (Ressler et al. 2011; Breen et al. 2018). Therefore, we investigated whether the vulnerability phenotypes detected here were associated with distinct epigenetic profiles. We assessed darters’ and freezers’ genome-wide transcriptional profiles in the ventromedial prefrontal cortex (vmPFC) (**Fig 5a-b**, **Supp. Table 1**), as this region is implicated in fear extinction, renewal, and PTSD (Milad and Quirk 2002; Kalisch et al. 2006; Peters et al. 2010; Stafford et al. 2012; Garfinkel et al. 2014; Marek et al. 2018).

**Figure 5.**
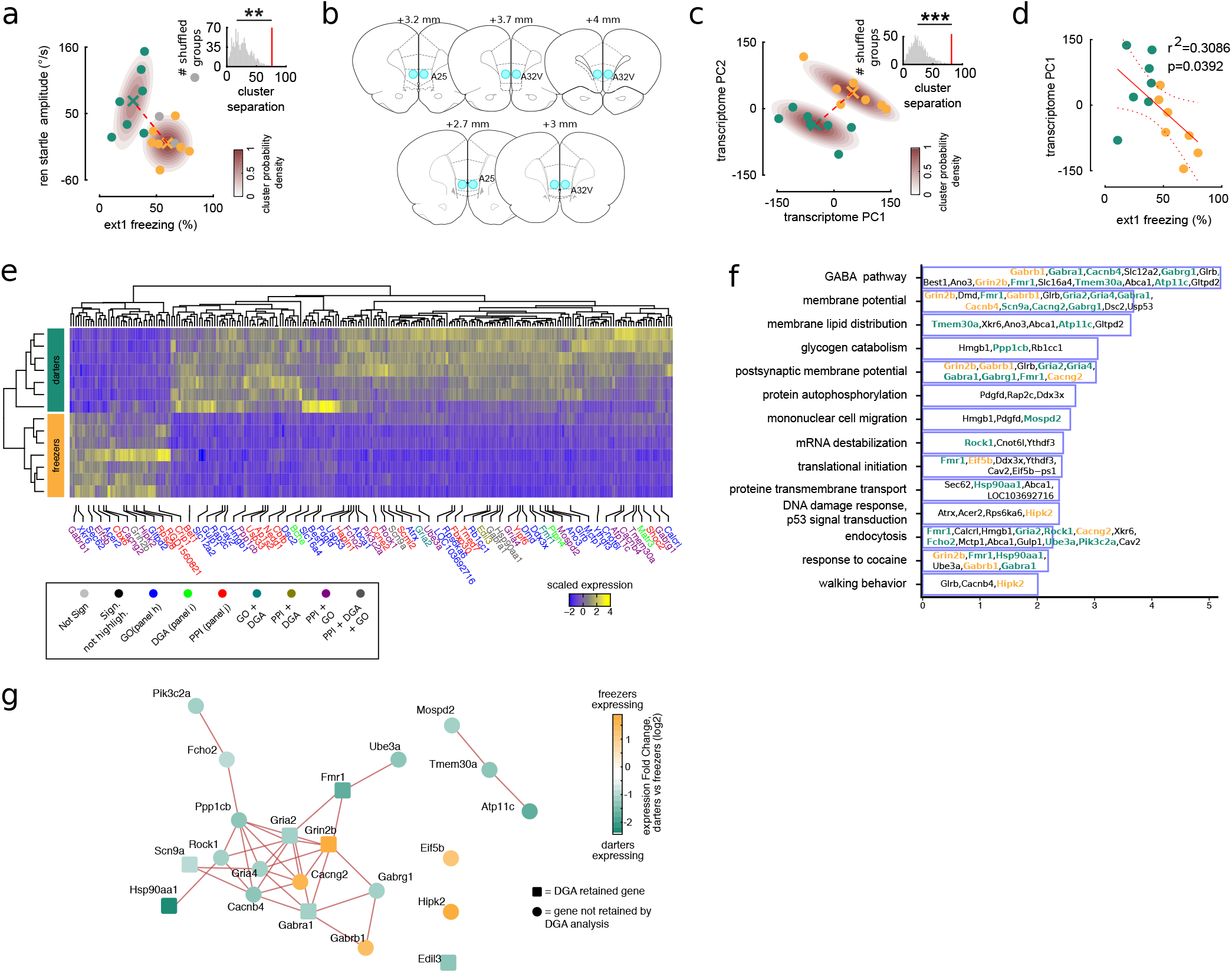
Darters and freezers have different gene expression profiles in ventromedial prefrontal cortex. (**a**) For transcriptomic analysis, two groups of equal size (n=7) were randomly subsampled from darters and freezers. Same as Fig. 4h, with gray dots representing the discarded data. (**b**) Brain coronal plane schema of the sampling zones (blue circles) for the transcriptomic analysis, corresponding to cingulate cortex areas 32V and 25. (**c**) Principal component analysis (PCA) of the vmPFC transcriptomes. The first two principal components represented 32% and 12% of the total variability in the data respectively. (**d**) Linear regression analysis of gene expression profile (represented by the first PCs of the transcriptome) vs. early extinction behavioral phenotypes. (**e**) Scaled expression of DEGs for darters and freezers. The dendrograms represent the hierarchical clustering of individuals and genes according to gene expression. Only the 78 genes selected by the GO (panel f), PPI (Supp. Fig. 6a), DGA (Supp. Fig. 6b) analyses are labeled. The colored lettering indicates those analyses for which the DEGs were retained. (**f**) Significantly enriched Biological Processes Gene Ontology (GO) terms. The genes in bold typeface correspond to those identified in at least two among GO, Protein-protein interaction (PPI, supp. Fig. 6a), and Disease-Gene Association (DGA, supp. Fig. 6b) analyses. Genes upregulated in freezers are in yellow while genes upregulated in darters are in green. (**g**) Summary of DEGs retained by at least two among GO, PPI, and DGA analyses. The intensity of the color of the node represents the relative expression of the corresponding gene in darters vs freezers. Lines represent the protein-protein associations.

Darters’ and freezers’ gene expression profiles clustered in two groups (**Fig 5c**), reflecting their behavioral phenotypes (**Fig. 5d**), and we found 238 genes expressed differentially (DEGs) between darters and freezers (**Fig. 5e**, **Supp. Fig. 5**, **Supp. Table 2**). Among those, a gene ontology (GO) analysis revealed significantly enriched pathways comprising 49 genes; the enriched pathways are involved in GABA signaling, regulation of membrane potential, membrane molecular organization, as well as glycogen catabolic processes (**Fig. 5f**, **Supp. Table 3**). In addition, we identified 45 DEGs involved in known protein-protein interaction (PPI) networks (**Supp. Fig. 6a**, **Supp. Table 4**). Finally, a disease-gene association (DGA) analysis revealed that 10 DEGs have previously been associated with PTSD, anxiety, or stress disorders (Roberts et al. 2016; Meier et al. 2019) (**Supp. Fig. 6b, Supp. Table 5**).

In some cases, these three analyses selected for the same DEGs. Indeed, 22 DEGs were highlighted by two or more analyses, marking them as possible key players in the vulnerability to fear relapse (**Supp. Fig. 6c**, **Fig. 5g**). Remarkably, these 22 genes were preferentially involved in synaptic functions and plasticity. In particular, higher fear renewal in darters was associated with an overexpression of GABAa signaling, while freezers, less vulnerable, overexpressed GABAb receptors and glutamate NMDA signaling receptors, required in vmPFC for fear extinction consolidation (Burgos-Robles et al. 2007). This is consistent with decreased GABAb signaling implication in fear extinction impairment as well as increased generalization of conditioned fear (Shaban et al. 2006; Lange et al. 2014; Zhang et al. 2016).

## DISCUSSION

Here, we showed that behavioral markers emerging during early fear extinction predict interindividual vulnerability to fear renewal and associated cortical gene expression profiles in male rats. These behavioral markers and genes may represent targets for novel treatments to enhance context generalization of extinction training.

Measuring freezing behavior in rodents in small conditioning chambers has been instrumental to study the neurobiological bases of fear and emotional processing. While freezing levels have been used as an index to study interindividual variability in fear behavior (Gruene et al. 2015; Dopfel et al. 2019), to date they do not provide clear biomarkers of the vulnerability to fear relapse after recovery. Outside the laboratory, both humans and rodents can express multiple defensive behaviors in response to a perceived threat, such as freezing, fleeing, or attacking (LeDoux and Daw 2018; Fanselow 2018). To improve the translation of findings from experiments with animal models to the clinic, one approach is to develop more naturalistic experiments to distinguish interindividual responses as well as the expression and measure of an enlarged possibility of behaviors (Mobbs and Kim 2015; Paré and Quirk 2017; Kim and Jung 2018; Headley et al. 2019). Consistently, we employed a new ethologically relevant paradigm combined with an automated pipeline for more extended behavioral profiling. This enabled the isolation of distinct behavioral phenotypes that could not be observed with standard spatially restricted behavioral settings which reduce natural behaviors such as foraging. Indeed, by showing that darting behavior during early extinction is a marker of fear relapse vulnerability, our results challenge the tenet that assessment of freezing alone is sufficient to model post-traumatic behavior in rodents. This novel behavioral paradigm may have direct applications in translational research, since it provides a direct way to identify subjects vulnerable to fear renewal early after the traumatic experience. For instance, translational research models may use this protocol to directly test the effectiveness of novel pharmacological strategies in reducing fear relapse vulnerability, and thus improve the treatment of patients suffering from anxiety disorders. It would be of interest for future studies to further characterize how darting vs. freezing propensity may depend upon the intensity of the shock since others have shown that increased foot shock intensity may decrease darting behavior (Mitchell et al. 2021). Nevertheless the darting observed in small environments is mostly characterized by jumping and may not be directly comparable to the darting response that we observed in our large open field arena, mainly expressed in running.

For the same reason, it is not directly possible to interpret our results in relation to previous work suggesting that defensive darting may be expressed to different degrees in males and females, with female rats displaying higher levels of darting (Gruene et al. 2015; Pellman et al. 2017; Colom-Lapetina et al. 2019; Greiner et al. 2019; Morena et al. 2021; Mitchell et al. 2021). Interestingly, here we observed darting in an all-male cohort of animals. This might be due to the larger dimensions of our open field compared to that of previous work. Further investigations testing female rats in our paradigm could provide a promising and needed perspective of this work.

Darting behavior could be interpreted either as a fearful defensive escape response or as an expression of locomotor behavior by a less frightened rat performing behaviors other than freezing. The CS-evoked darting we observed in the large open field may be related to fear-related conditioned jumping previously observed in small conditioning chambers (Gruene et al. 2015; Fadok et al. 2017; Totty et al. 2021), which likely corresponds to attempted escape reponses. Consistent with this, darting was triggered by the CS onset, decreased with extinction training, and darting trajectories had the tendency to start and end in safer spots of the environment, all consistent with darting as a CS-evoked defensive behavior. Future work should investigate whether darting behavior might be assimilated to escape responses, and where it would figure in the continuum of fear states. The transition between darting and freezing may depend upon the characteristics and perceived level of safety of the current environment, or upon the physical and psychological distance from the perceived threat (Blanchard et al. 1986; Fanselow and Lester 1988). In this view, CS-evoked darting in the open field would be interpreted as a defensive response of animals perceiving the threat as being less imminent than it was perceived by the freezers. Therefore, these animals would be expected to be less vulnerable towards post-extinction relapse and freeze less. Surprisingly, we showed the opposite, indicating that freezing as an index of fear does not transfer proportionally across environments. Perhaps, the darters expressed greater fear renewal in the form of freezing and startle because the constrained environment rendered their preferred defense, darting responses, impossible. Future studies could explore this by testing renewal in an environment allowing darting but sufficiently different from the extinction environment to induce a renewal effect.

Vulnerability to fear renewal might also be driven by differential processing of contextual information (Maren et al. 2013), since the darters had a higher context dependence of extinction, expressed in the form of higher fear renewal. Possibly, darters are less frightened than freezers during extinction in the open field, and this lower fear expression may indicate lower attention levels to the CS presentation and therefore poorer extinction learning than freezers. Weaker extinction learning would thus explain why darters are more vulnerable to fear renewal. However, the fact that within-session extinction learning rates are not different between freezers and darters argues against this interpretation. Nevertheless, long-term extinction learning efficacy across days could rely also on post-training consolidation mechanisms (Datta and O’Malley 2013), and therefore may not be reflected by within-session learning rates.

Identifying and understanding the biological mechanisms underlying interindividual responses to trauma and therapy is crucial for effective personalized therapy for patients. Here, we made a step towards deciphering the transcriptional coding landscape that is specifically associated with interindividual fear renewal vulnerability. We found that DEGs between more and less vulnerable subjects included genes involved in GABA signaling pathways, as well as in the regulation of membrane potential and membrane proteic organization. These might be crucial for individual vulnerability to fear relapse by regulating long term synaptic plasticity mechanisms in the vmPFC, similarly to what has been reported in the amygdala for fear conditioning (Shaban et al. 2006; Lange et al. 2014). Our results provide additional support of current clinical trials showing that the pharmacological facilitation of NMDA receptors activity might augment EBT efficacy, at least in some patients (Mataix-Cols et al. 2017). The observed transcriptional changes could be regulated by DNA (Soliman et al. 2010; Stafford et al. 2012) and RNA epigenetic mechanisms (Lin et al. 2011; Li et al. 2019). Future studies should focus on changes in these regulatory pathways through conditioning, extinction, and renewal in order to better understand their respective roles in interindividual vulnerability. In addition, the study of behavioral phenotype related gene expression profiles within specific neuron types or populations (Chen et al. 2020) in vmPFC might be instrumental to understanding the cellular specificity of the implicated pathways.

## METHODS

For detailed information about materials and methods, see the Supplement.

### Behavioral data acquisition

18 adult male Long-Evans rats received a surgical implantation of a magnetic base onto which was fixed the Inertial Measurements Unit (IMU) during the experiments. The IMU is a small and lightweight device containing accelerometers and gyroscopes that sample linear acceleration and angular velocity of the animals’ head in three dimensions (Pasquet et al. 2016).

After recovery from surgery, the rats underwent a 14 day ABC fear renewal protocol (Bouton 2004), which employed three different environments: two standard fear conditioning chambers (A and C) and one open field arena B. Auditory cues (CS) were 20 s continuous pure tones at 2 kHz (62-68 dB). Each day the rat had one training/testing session. Cameras mounted on the sides of the environments monitored animals’ behaviors while a ceiling mounted video tracked their positions. During habituation (days 1 through 5), animals were habituated to the open field during 26 min sessions of free exploration and foraging. During fear conditioning (days 6 and 7), rats underwent two fear conditioning sessions in A, each composed of a 10 min baseline recording followed by five presentations of the CS each co-terminating with a footshock (1 s; 0.7 mA) at 10 minute intervals. Rats were then left undisturbed in their cages during days 8 and 9, during the weekend. Extinction: on days 10 to 13, rats underwent extinction training in B. Each session consisted of a 6-minute baseline recording followed by three presentations of the CS at intervals of 6 minutes. Fear renewal test: on day 15, all rats underwent a fear renewal test in C. After 6 min baseline recording, three CS were presented at 6 minute intervals.

### Behavioral data analysis

Freezing was defined as each continuous period when the angular speed, computed from the IMU gyroscopes, was below 12°/s for at least 200 ms, as shown previously (Pasquet et al. 2016). Since the animals were not equipped with the IMU during conditioning, freezing was manually scored for these sessions, and was defined as the absence of movement except for breathing. Object exploration was estimated as the time the animals spent within 5 cm around the objects. To detect darting, LED position data was smoothed with a 300 ms Gaussian window. Then, we first identified intervals when the animal moved at least 15 cm without changing direction (IMU-detected angular speed inferior to 3°/s in bins of 50 ms). The resulting straight trajectories were divided into darting vs. slower movement epochs by a k-means classification on the animal’s speed. A supervised deep learning algorithm scored the remaining behaviors. To create the training and test data sets, an experienced experimenter manually scored behaviors into the following categories: two types of grooming (face and body), rearing, freezing, and darting. The average accuracy of the classification was 87.7% (Supp. Fig. 7a), consistent with previous reports (Venkatraman et al. 2010).

Using all data over the course of the entire protocol for all sessions of all the animals, a six-column matrix containing the time series of the respective classified behaviors was built and smoothed (Gaussian window of 20 s). All of the CS and post-CS (60 s) data from the five extinction sessions of all of the animals was extracted and its dimensionality was reduced with Principal Component Analysis (PCA). Then, unsupervised clustering (k-means) separated the animals into two groups using the average activation strengths of the first three components of the PCA during the three CS presentations of the first extinction session (ext1). As a control, data was k-means clustered into two groups from data from the three CS presentations of the last session of extinction (ext5). Behavioral analyses were focused on CS evoked responses taking place in the intervals from CS onsets until 60 seconds after CS offsets, here named CS periods.

### Transcriptomic analysis

Following the renewal protocol, rats were euthanized and their brains immediately removed and frozen. RNA was extracted from micro-punch of the ventromedial prefrontal cortex (vmPFC). Genomewide transcriptional profiling was performed for seven rats in each group. The analyses were performed using the Eoulsan pipeline (Jourdren et al. 2012), including read filtering, mapping, alignment filtering, read quantification, normalisation and differential analysis. Before mapping, poly N read tails were trimmed, reads ≤40 bases were removed, and reads with quality mean ≤30 were discarded. All overlapping regions between alignments and referenced exons were counted using HTSeq-count 0.5.3 (Anders et al. 2015). Statistical treatments and differential analyses were performed using DESeq2 1.8.1. Differentially expressed genes (DEGs) were defined by an absolute value of log2 of fold change > 1, an adjusted pvalues after Benjamini & Hochberg correction (Benjamini and Hochberg 1995) of <0.01, and mean total counts for all the conditions > 10. Gene annotation and enrichment analysis on the DEGs was performed using Metascape (Zhou et al. 2019). Protein-protein interactions analysis between the DEGs was performed using STRING (Search Tool for Recurring Instances of Neighboring Genes, v 11.0; Szklarczyk et al. 2016). Disease Gene association between the DEGs and the stress and anxiety-related disorders was performed using DisGeNET database (Piñero et al. 2020).

## ACKNOWLEDGMENTS

We thank Matthieu Pasquet and Guillaume Dugué for help with inertial data acquisition and analysis, Karine Dias and Sophie Lemoine for RNA sequencing and raw transcriptomic data postprocessing, Cédric Colas, Benjamin Billot, and Pierre-Antoine Vigneron for help with setting up of the apparati and protocols, and Valérie Doyère, Michaël Zugaro, and Gabrielle Girardeau for useful comments on the manuscript. This project was funded with grants from the Agence Nationale de la Recherche to BPG, TMJ, and SIW, Labex MemoLife to MNP and SIW, Fondation Pierre Deniker to MNP, and Fondation de France to M-OK. FD was supported by the Fondation pour la Recherche Medicale.

## AUTHOR CONTRIBUTIONS

Research project conception and design: MNP, FD, BPG, TMJ. Funding gathering: BPG, TMJ, SIW, MNP, M-OK, FD. Technological implementation: MNP. Experiments: FD, MNP. Data analysis design: MNP, FD, RT, SIW, GM. Data analysis implementation: FD, MNP, RT, GM, AF. Visualization: MNP. Manuscript: MNP, FD, SIW, RT with comments from all authors. Project supervision and management: MNP.

## COMPETING INTERESTS

The authors declare no competing interests.

## SUPPLEMENTARY MATERIALS

### SUPPLEMENTARY FIGURES

**Supplementary Figure 1.**
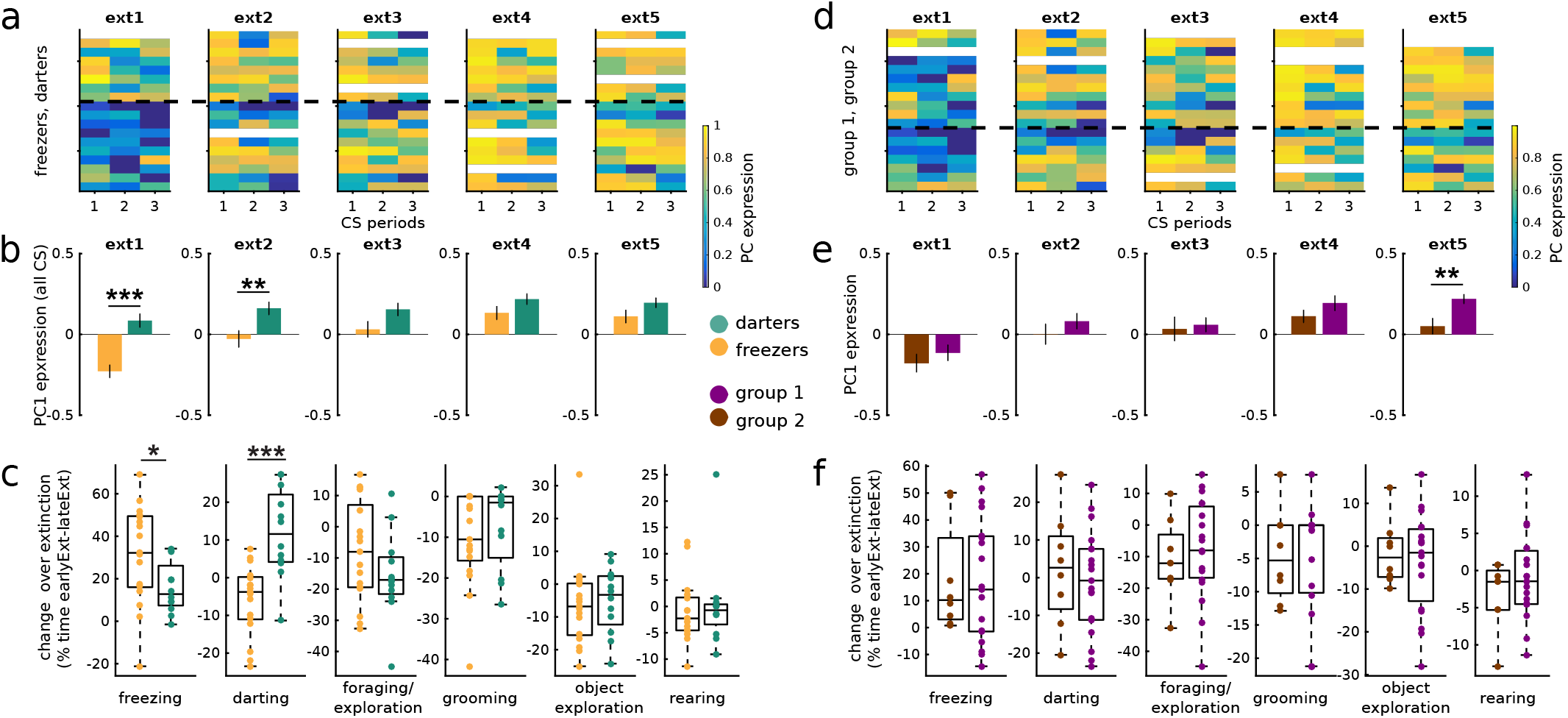
Segregating animals based on behavioral phenotype data from ext1 (a-c) but not from ext5 (d-f) leads to groups with significant differences of behavior evolution over extinction training. (**a**) Evolution of PC1 values over the three CS periods in freezer and darter groups as clustered from ext1 data. Each row corresponds to one rat (white corresponds to unavailable data). Colors represent the average expression of PC1 for the respective CS presentations (numbered below). The horizontal dashed line separates the two groups (freezers and darters) determined from ext1 data. (**b**) Average values of PC1 expression over the sessions for the two groups. The expression of PC1 is significantly different between the two groups in ext1 (as expected) and also ext2 (unpaired t-test; **p<0.01, ***p<0.001). (**c**) Changes in the behavior expression across extinction training sessions for the two groups as clustered with ext1 data. Colored dots represent the difference between early extinction (average of ext1 and ext2) and late extinction (average of ext4 and ext5). Note how the two groups manifest different evolution of the expression of freezing and darting over extinction training: darting extinguishes in darters and freezing change over extinction is greater for freezers. (**d**,**e**,**f**) Same as (a,b,c) but for the groups derived from k-means clustering on ext5 data. Note that the expression of PC1 differs between the two control groups only for the ext5 session used for clustering. Clustering with ext5 data did not lead to significant inter-group differences in behavioral expression changes over extinction. Data is represented as mean ±SEM. Box plot format is the same as in Fig 1. Statistics are unpaired t-tests (*p<0.05, **p<0.01, ***p<0.001).

**Supplementary Figure 2.**
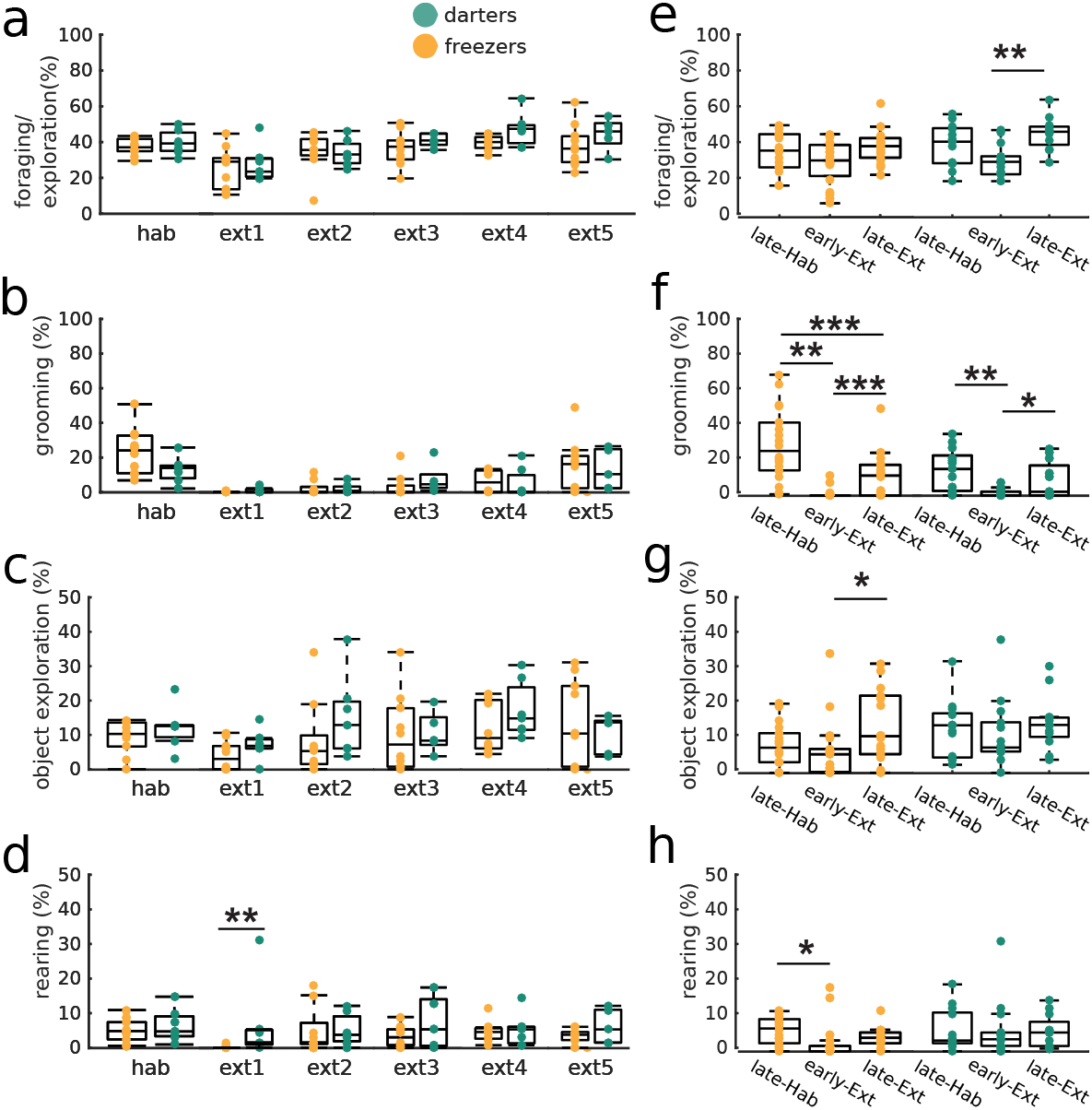
Comparison of expression of behaviors other than darting and freezing during CS periods across extinction training. (a-d) Equivalent of Figure 3a-b for other behaviors. (e-h) Intra-group comparisons demonstrate significant differences in these behaviors expression over extinction learning equivalent of Figure 3c-d for other behaviors. Statistics are unpaired t-tests (*p<0.05, **p<0.01, ***p<0.001)

**Supplementary Figure 3.**
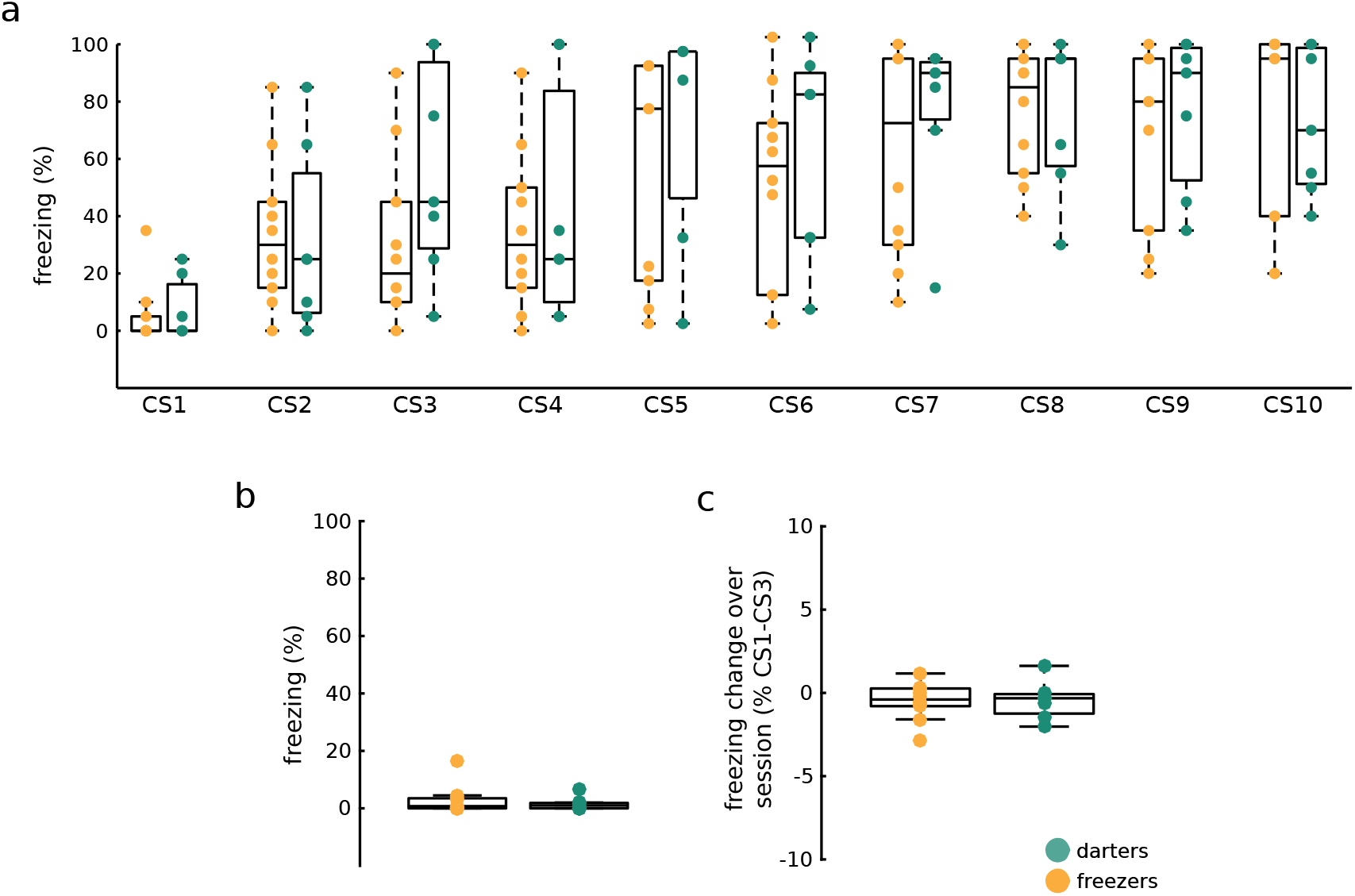
Fear acquisition, generalization of contextual fear, and within-day learning rate do not explain interindividual variability during early extinction. (**a**) Percentage of time spent freezing during CS presentations in conditioning sessions (CS1-5 were presented on day 1 and CS6-10 on day 2). (**b**) Percentage of time the animals spent freezing in baseline period prior to the first CS presentation in ext1. (**c**) Differences between CS3 and CS1 for proportion of time spent freezing in ext1. No significant differences (Rank-sum tests p>0.05). Box plot format is the same as in Fig 1.

**Supplementary Figure 4.**
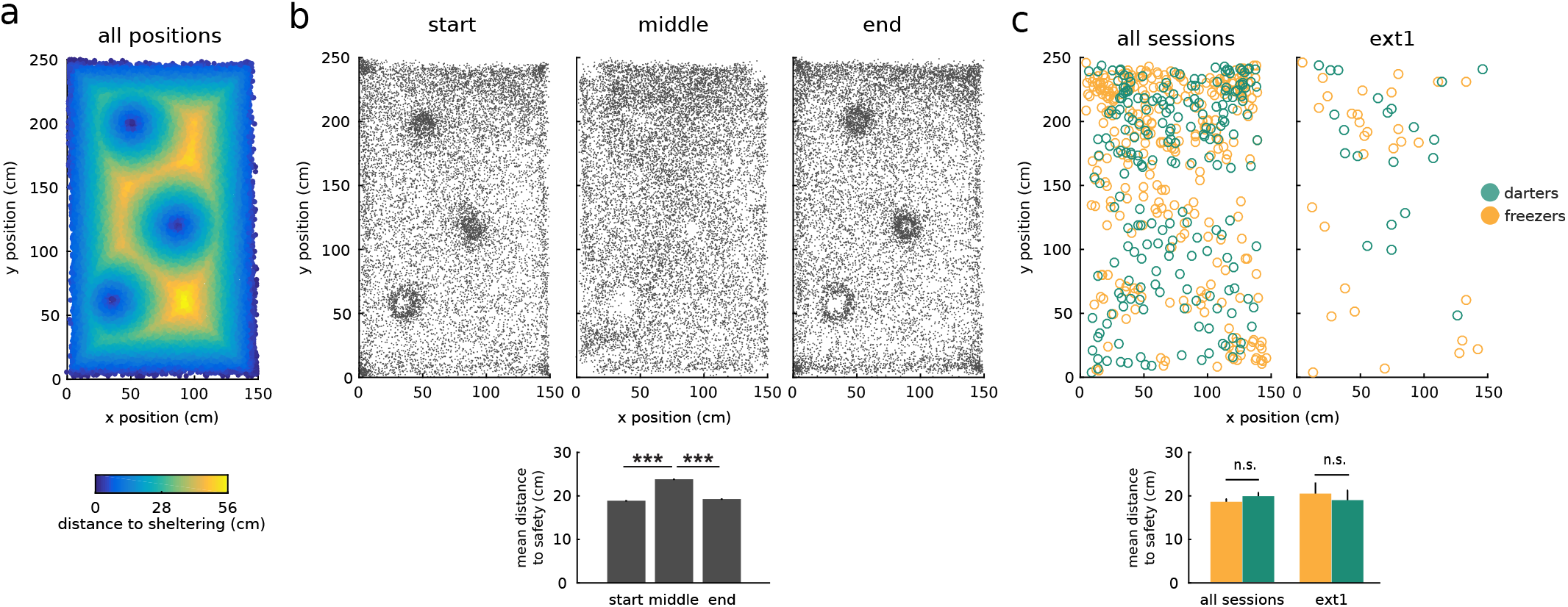
Darting typically was from one ‘sheltering’ location to another. (**a**) Positions in the open field are colored according to their distance from the walls and the objects (all considered as ‘sheltering’ locations). (**b**) Dots indicate all rats’ positions at the beginning (top left), in the middle (top middle), and at the end (top right) of all darting trajectories for all sessions in one configuration of the open field (see Methods). (bottom) When in the middle of darting trajectories, rats were further away from sheltering locations (‘safety’) than at the beginning or end of trajectories. (**c**) Positions of all rats (color coded by group) at CS onset for all sessions (left) and for ext1 (right). (bottom) No significant distance-to-safety difference between groups at CS onset. Data represented as mean ±SEM. Unpaired t-tests. [n.s.:not significant; ***p<0.001.]

**Supplementary Figure 5.**
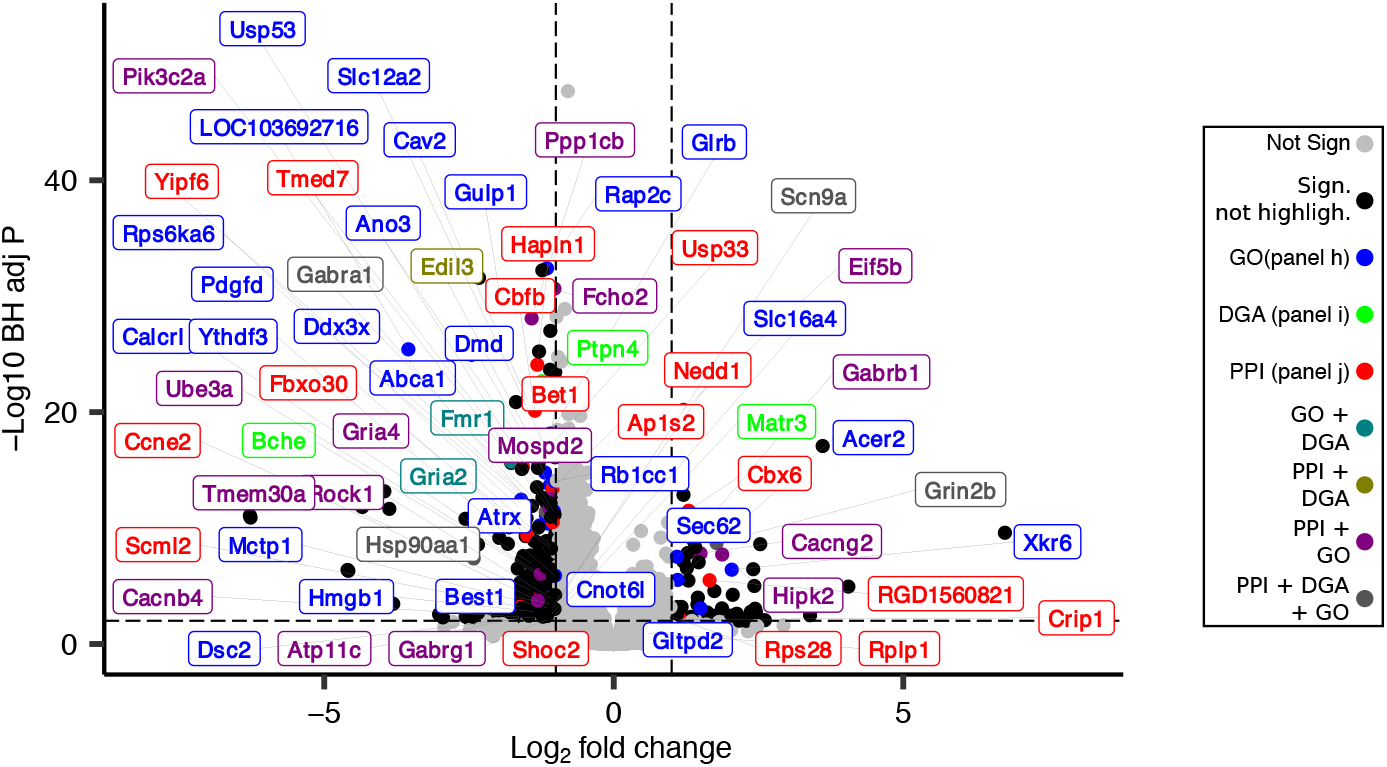
Volcano plot of the differential expression of the genes. Genes were considered to be significantly differentially expressed between groups if their adjusted P (Padj) was <0.01 (horizontal dotted line) and the absolute value of their Fold Change was >2 (vertical dotted lines). Non-DEGs are depicted by light gray dots. The colored lettering indicates DEGs retained by GO, PPI, and DGA analyses.

**Supplementary Figure 6.**
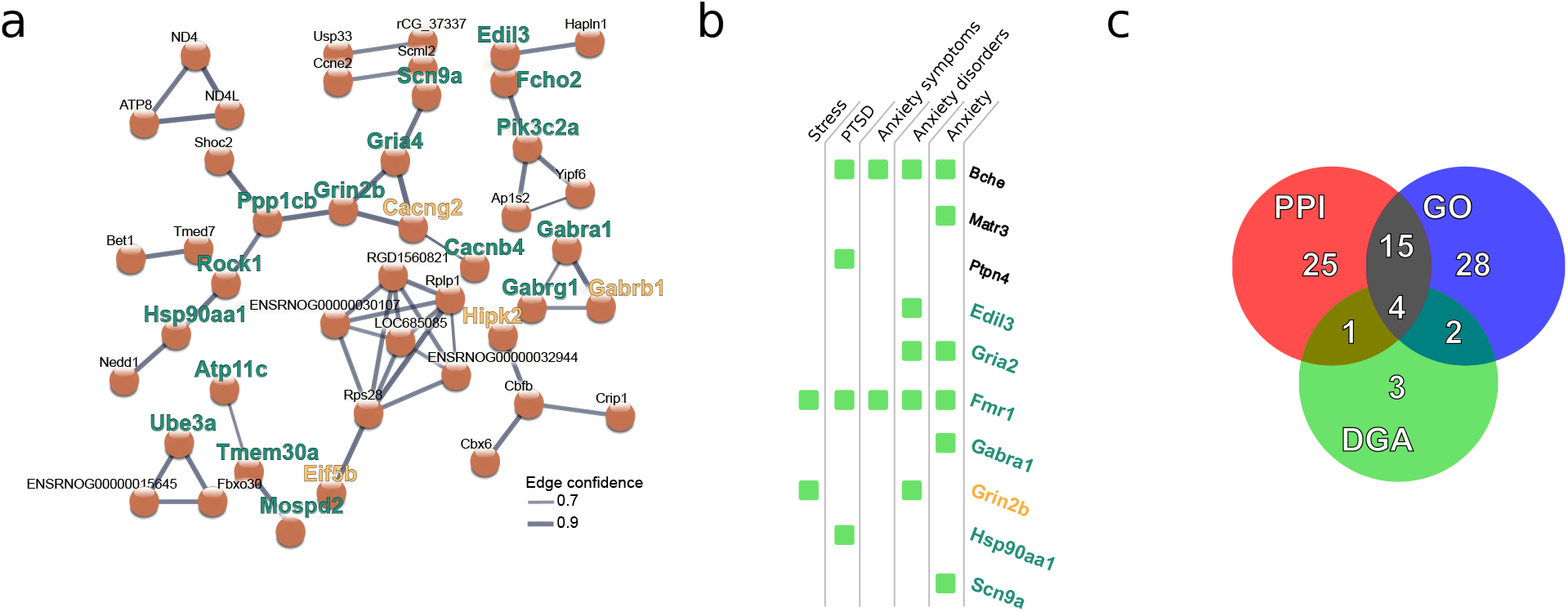
Selection of DEGs more likely to play a role in the vulnerability to fear relapse (**a**) PPI network of the DEGs obtained from STRING analysis of the DEGs list. Line thickness represents the confidence level supporting each protein–protein associations. Only connected nodes and interactions with confidence >0.7 are shown. The network has significantly more interactions than expected with a PPI enrichment p-value ꜟ0.02. Genes upregulated in freezers are in yellow while genes upregulated in darters are in green. (**b**) DGA analysis using the DisGeNET database revealed DEGs previously associated with anxiety and stress-related disorders, indicated by the green squares. Gene label color-coding as in Figure 6. (**c**) Venn diagram of the categories of the 78 DEGs selected by the GO Terms, PPI and DGA analyses.

**Supplementary Figure 7.**
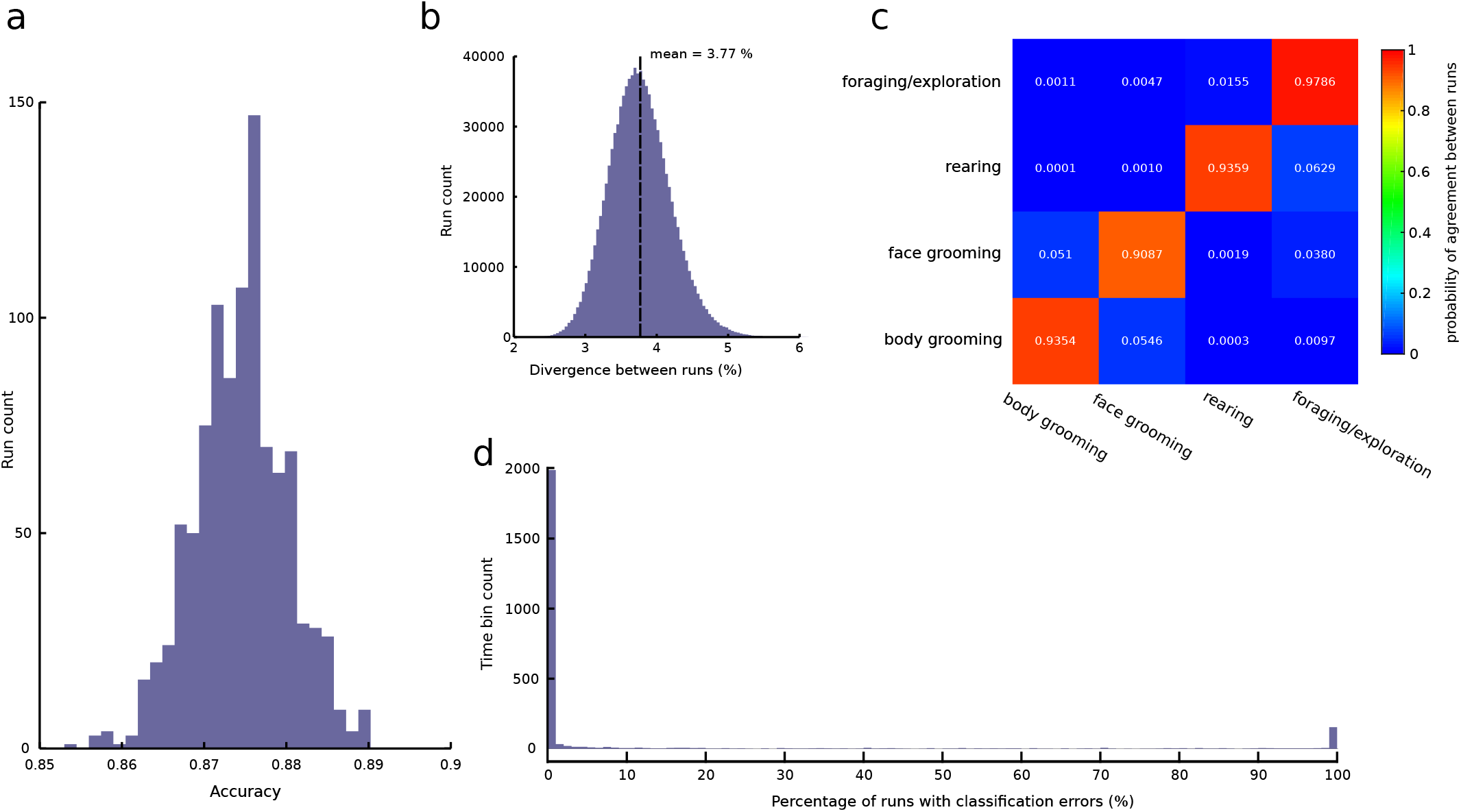
Performance and outcome consistency of supervised machine learning classification of behaviors from IMU and camera data over 1000 runs (see Methods). (**a**) Classification accuracy distribution. (**b**) Percent divergence between different runs of the machine learning classification. Results of the different runs were compared to each other, and for each comparison a divergence value was computed corresponding to the percentage of time bins with different classifications. On average, the different runs were different in 3.77% of the bins and the divergence was never greater than 6%. (**c**) Conditional probability of classification. Colors represent the probability (also expressed numerically for each cell) that one run classified a time bin to a particular behavior, given the classification outcome for that bin from another run. For example, the top left cell indicates that the bins that were classified as “foraging/exploration” on one run, had 0.001 (or 0.1%) probability of being classified as “body grooming” on another run. Note the high values along the diagonal, indicating that time bins classified as a particular behavior on a given run are very likely to be classified as the same behavior on a different run, and therefore demonstrating consistency between runs. (**d**) For each of the 2514 time bins of the test dataset, the proportion of runs when an error occurred was computed. Errors tend to concentrate in a few time bins and most time bins were never wrongly classified.

## SUPPLEMENTARY VIDEOS

**Supplementary Video 1** Examples of the 6 behavioral categories.

The video is provided as a separate file.

**Supplementary Video 2** Example of a CS-evoked orienting response and associated startling in a small compartment.

The video is provided as a separate file.

## SUPPLEMENTARY TABLES

**Supplementary table 1**. Summary of RNAseq results. Results of the 32623 analyzed genes and corresponding statistics are detailed.

The table is provided as a separate spreadsheet.

**Supplementary table 2.**
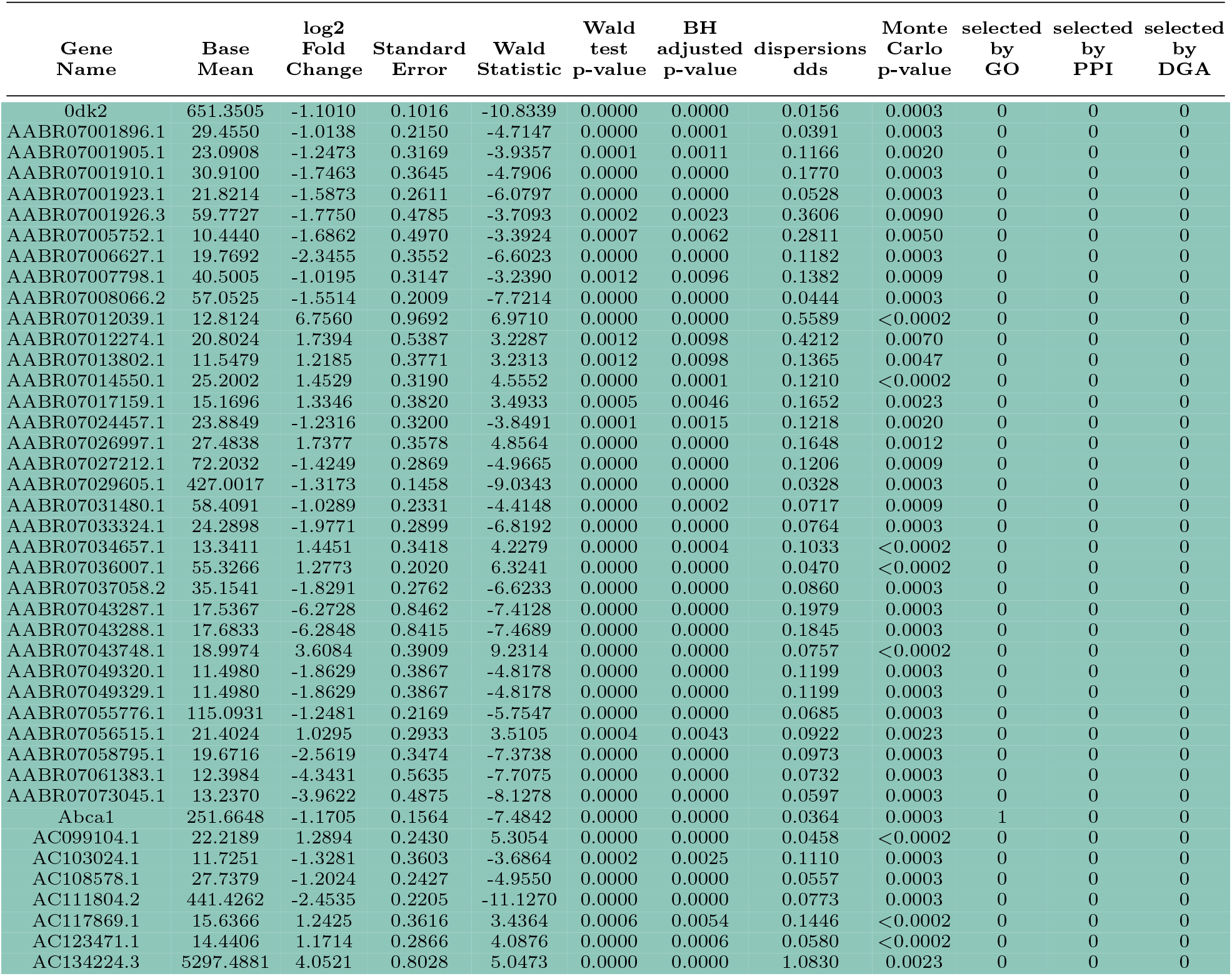
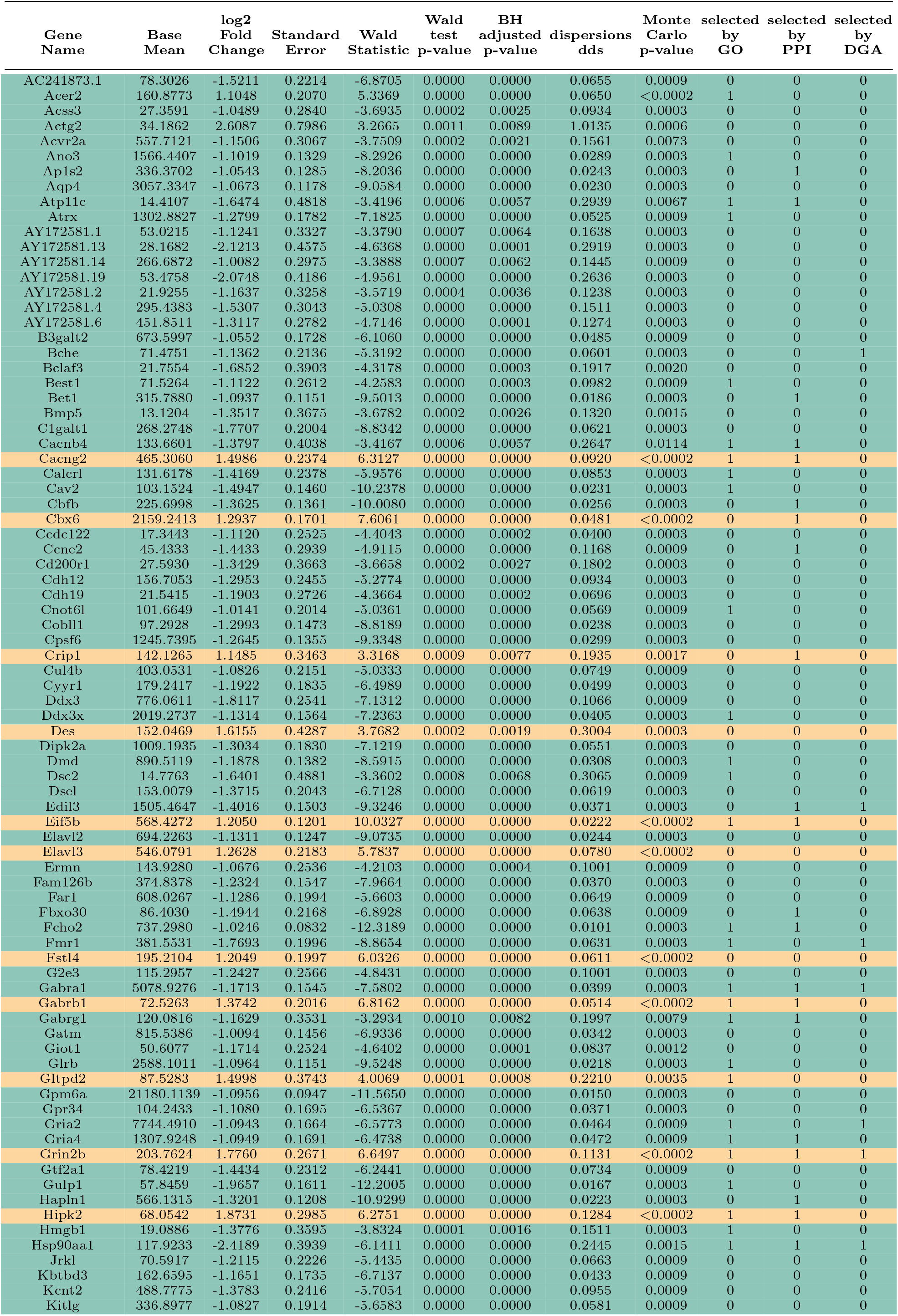
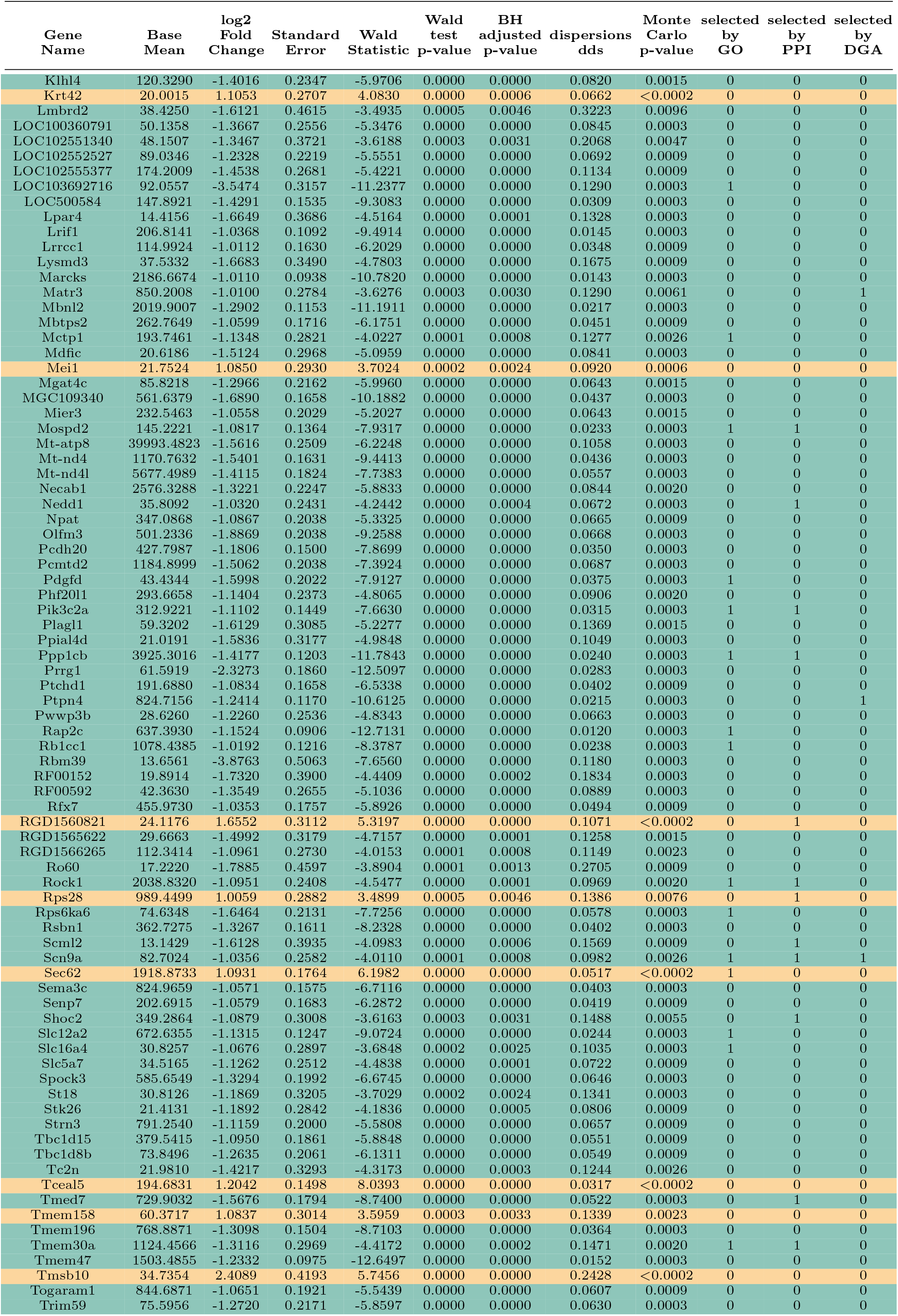
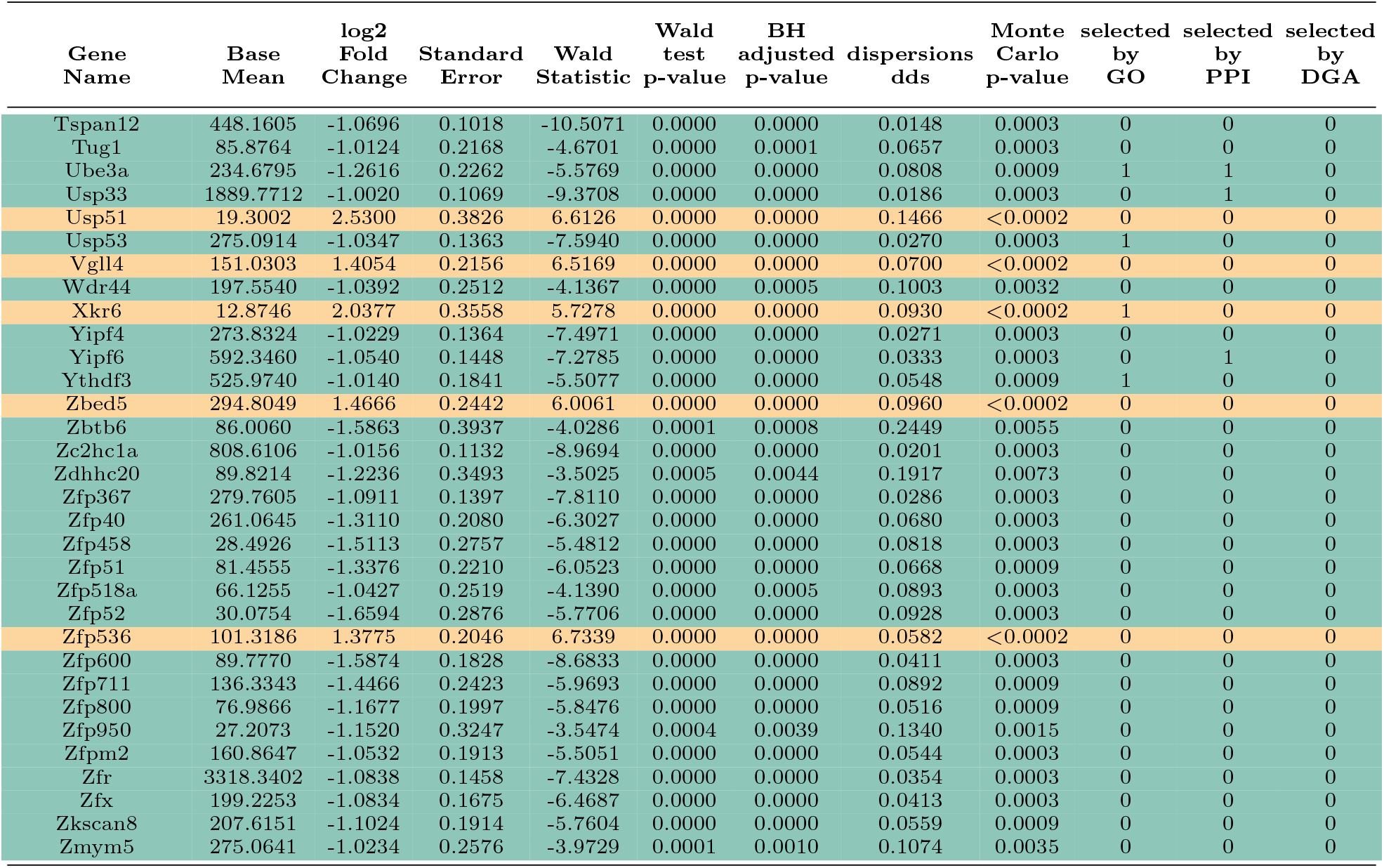
Summary of all Differentially Expressed Genes (DEGs) and corresponding statistics found in the genes listed in Supplementary table 1. Genes are ordered alphabetically and corresponding rows are colored in green or yellow for genes expressed more in darters or freezers, respectively. In the last three columns the number 1 indicates that the corresponding gene was selected by GO, PPI, and DGA analyses.

**Supplementary table 3.**
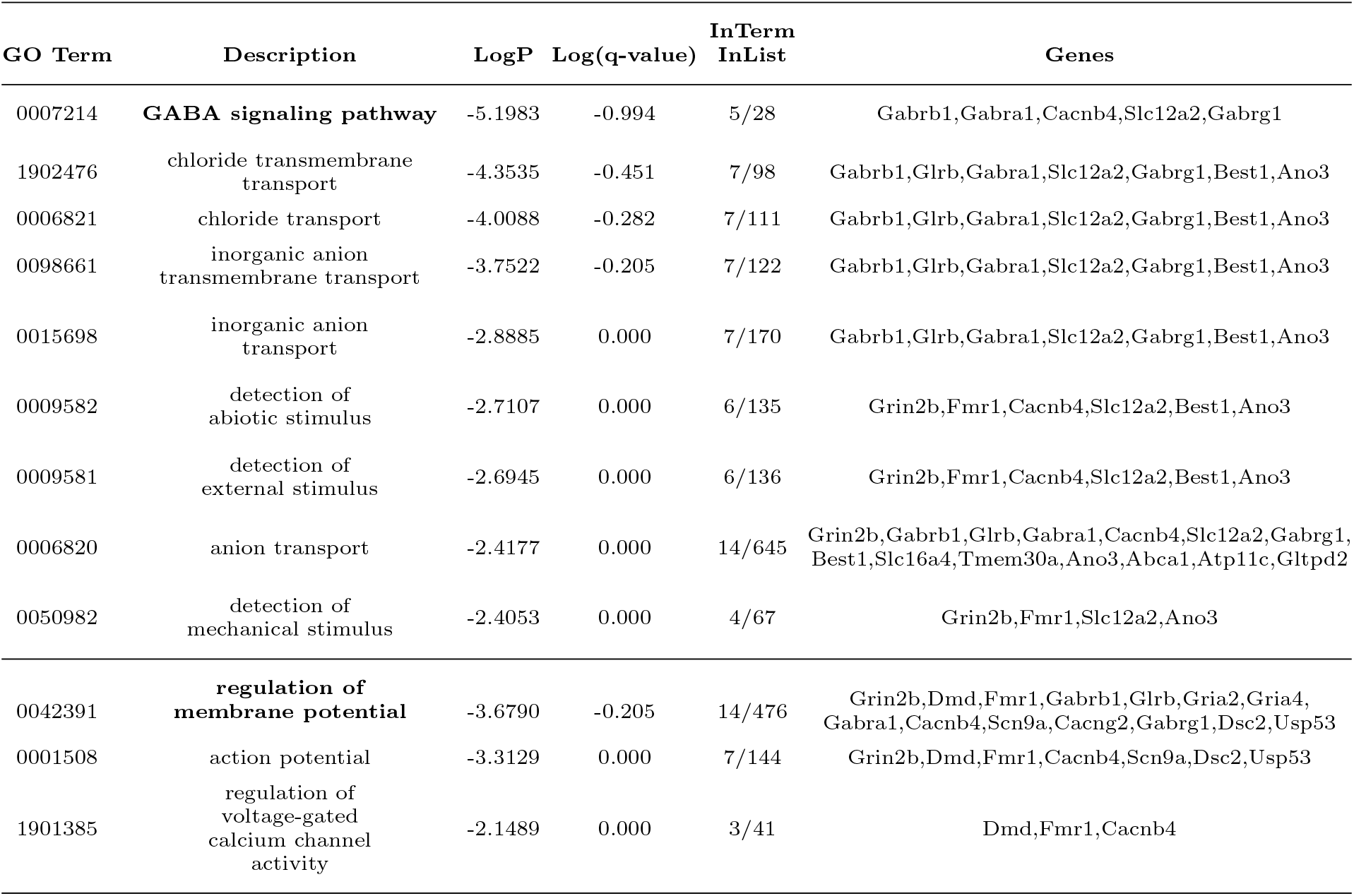
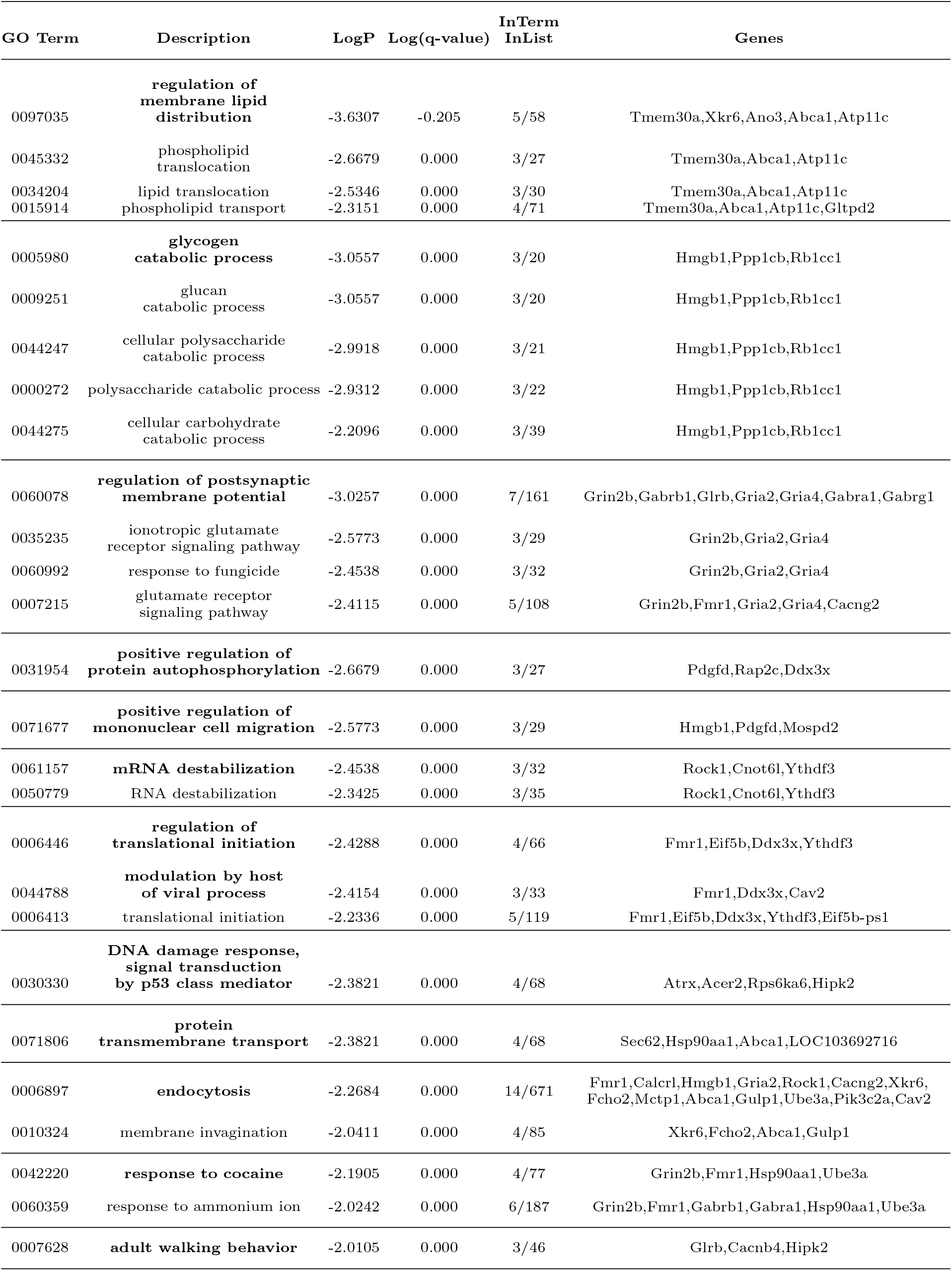
Summary of GO analysis results. Significantly enriched Biological Processes GO terms are represented. P-values were calculated based on the accumulative hypergeometric distribution, and q-values were calculated using the Benjamini-Hochberg procedure to account for multiple testing using Metascape. The portion of genes associated with each GO term that was identified within the DEGs are reported and their symbols are detailed. Horizontal lines separate groups of GO terms whose summary term is indicated in bold.

**Supplementary table 4.** STRING analysis summary and statistics. Each protein-protein interaction that is part of the PPI network of the DEGs obtained from STRING analysis of the DEGs list is described. Only connected nodes and interactions with confidence >0.7 are described. The strength of data supporting each association is reported as well as the type of connection: Neighborhood on chromosome, Homology, Co-expression, Experimentally determined interaction, Analysis of database information, Automated text mining of co-occurrence of gene/protein names.

| Node 1 | Node 2 | Neighborhood<br>on chromosome | Homology | Coexpression | Experimentally<br>determined<br>interaction | Database<br>annotated | Automated<br>textmining | Combined<br>score |
| --- | --- | --- | --- | --- | --- | --- | --- | --- |
| ATP8 | ND4L | 0 | 0 | 0.718 | 0 | 0 | 0.878 | 0.964 |
| ATP8 | ND4 | 0 | 0 | 0.248 | 0 | 0 | 0.860 | 0.890 |
| Ap1s2 | Yipf6 | 0 | 0 | 0.083 | 0 | 0.900 | 0.049 | 0.905 |
| Ap1s2 | Pik3c2a | 0 | 0 | 0.045 | 0 | 0.900 | 0.043 | 0.900 |
| Atp11c | Tmem30a | 0 | 0 | 0.087 | 0.222 | 0 | 0.664 | 0.741 |
| Bet1 | Tmed7 | 0 | 0 | 0.110 | 0 | 0.900 | 0.078 | 0.910 |
| Cacnb4 | Cacng2 | 0 | 0 | 0.098 | 0 | 0.900 | 0.273 | 0.928 |
| Cacng2 | Grin2b | 0 | 0 | 0.167 | 0.127 | 0.900 | 0.584 | 0.965 |
| Cacng2 | Gria4 | 0 | 0 | 0.126 | 0.403 | 0.900 | 0.838 | 0.990 |
| Cbfb | Crip1 | 0 | 0 | 0.063 | 0.806 | 0 | 0 | 0.810 |
| Cbfb | Hipk2 | 0 | 0 | 0 | 0 | 0.900 | 0.058 | 0.901 |
| Cbfb | Cbx6 | 0 | 0 | 0.062 | 0 | 0.900 | 0 | 0.902 |
| Ccne2 | Scml2 | 0 | 0 | 0.064 | 0.780 | 0 | 0 | 0.785 |
| ENSRNOG<br>00000015645 | Fbxo30 | 0 | 0 | 0.111 | 0 | 0.900 | 0 | 0.907 |
| ENSRNOG<br>00000015645 | Ube3a | 0 | 0 | 0.063 | 0.051 | 0.900 | 0.065 | 0.905 |
| ENSRNOG<br>00000030107 | Rplp1 | 0 | 0 | 0.540 | 0.615 | 0 | 0.151 | 0.836 |
| ENSRNOG<br>00000030107 | LOC685085 | 0 | 0 | 0.468 | 0.489 | 0 | 0.075 | 0.726 |
| ENSRNOG<br>00000030107 | RGD1560821 | 0 | 0 | 0.532 | 0.605 | 0 | 0.126 | 0.824 |
| ENSRNOG<br>00000030107 | Rps28 | 0 | 0 | 0.526 | 0.606 | 0 | 0.191 | 0.835 |
| ENSRNOG<br>00000032944 | Rplp1 | 0 | 0 | 0.540 | 0.615 | 0 | 0.151 | 0.836 |
| ENSRNOG<br>00000032944 | Rps28 | 0 | 0 | 0.526 | 0.606 | 0 | 0.191 | 0.835 |
| ENSRNOG<br>00000032944 | LOC685085 | 0 | 0 | 0.468 | 0.489 | 0 | 0.075 | 0.726 |
| ENSRNOG<br>00000032944 | RGD1560821 | 0 | 0 | 0.532 | 0.605 | 0 | 0.126 | 0.824 |
| Edil3 | Hapln1 | 0 | 0 | 0.108 | 0 | 0 | 0.834 | 0.846 |
| Eif5b | Rps28 | 0.092 | 0 | 0.081 | 0.917 | 0 | 0.126 | 0.931 |
| Fbxo30 | Ube3a | 0 | 0 | 0.064 | 0 | 0.900 | 0 | 0.902 |
| Fcho2 | Pik3c2a | 0 | 0 | 0.087 | 0 | 0.900 | 0.059 | 0.906 |
| Gabra1 | Gabrb1 | 0 | 0.813 | 0.223 | 0.649 | 0.900 | 0.761 | 0.974 |
| Gabra1 | Gabrg1 | 0 | 0.911 | 0.271 | 0.215 | 0.540 | 0.661 | 0.730 |
| Gabrb1 | Gabrg1 | 0 | 0.844 | 0.226 | 0.396 | 0.540 | 0.904 | 0.799 |
| Gria4 | Scn9a | 0 | 0 | 0.140 | 0.066 | 0.900 | 0.190 | 0.926 |
| Gria4 | Grin2b | 0 | 0.700 | 0.221 | 0.140 | 0.900 | 0.696 | 0.942 |
| Grin2b | Ppp1cb | 0 | 0 | 0 | 0.083 | 0.800 | 0.103 | 0.821 |
| Hsp90aa1 | Nedd1 | 0 | 0 | 0 | 0 | 0.900 | 0 | 0.900 |
| Hsp90aa1 | Rock1 | 0 | 0 | 0.080 | 0.176 | 0.900 | 0.118 | 0.924 |
| LOC685085 | Rplp1 | 0 | 0 | 0.543 | 0.615 | 0 | 0.130 | 0.833 |
| LOC685085 | Rps28 | 0 | 0 | 0.526 | 0.605 | 0 | 0.213 | 0.839 |
| LOC685085 | RGD1560821 | 0 | 0 | 0.534 | 0.605 | 0 | 0.106 | 0.821 |
| Mospd2 | Tmem30a | 0 | 0 | 0.080 | 0.144 | 0.900 | 0 | 0.914 |
| ND4 | ND4L | 0 | 0 | 0.799 | 0.871 | 0.360 | 0.955 | 0.999 |
| Pik3c2a | Yipf6 | 0 | 0 | 0.064 | 0 | 0.900 | 0 | 0.902 |
| Ppp1cb | Rock1 | 0.042 | 0 | 0.086 | 0.163 | 0.800 | 0.112 | 0.846 |
| Ppp1cb | Shoc2 | 0 | 0 | 0.064 | 0.238 | 0.720 | 0.422 | 0.869 |
| RGD1560821 | Rplp1 | 0 | 0 | 0.571 | 0.742 | 0 | 0.101 | 0.891 |
| RGD1560821 | Rps28 | 0.088 | 0 | 0.558 | 0.645 | 0 | 0.128 | 0.858 |
| Rplp1 | Rps28 | 0 | 0 | 0.855 | 0.742 | 0 | 0.327 | 0.972 |
| Usp33 | rCG_37337 | 0 | 0 | 0.799 | 0 | 0 | 0.067 | 0.805 |

**Supplementary table 5.** Summary of DGA analysis results and statistics. List of DEGs that have been previously associated with anxiety and stress-related disorders according to DGA analysis (DisGeNET database). The Disease Specificity Index (DSI) reflects whether genes are associated to several or fewer diseases and the Disease Pleiotropy Index (DPI) reveals whether these diseases are similar or not. The probability of being loss-of-function (LoF) intolerant (pLI) measures how much the naturally occurring LoF variation has been depleted from a gene by natural selection. LoF intolerant genes will have a high pLI value (>0.9). The GDA score indicates the level of evidence of the association based on the number and type of sources.

| Disease | Disease id | Gene | Gene Name | Disease Specificity Index | Disease Pleiotropy Index | pLI | GDA Score | # PMIDs | First PMID | Last PMID |
| --- | --- | --- | --- | --- | --- | --- | --- | --- | --- | --- |
| Anxiety | C0003467 | BCHE | butyrylcholinesterase | 0.447 | 0.923 | 1.06E-13 | 0.05 | 5 | 2017 | 2019 |
| Anxiety | C0003467 | FMR1 | FMRP translational regulator 1 | 0.473 | 0.769 | 0.64718 | 0.2 | 11 | 2002 | 2019 |
| Anxiety | C0003467 | GABRA1 | GABA A receptor subunit alpha1 | 0.563 | 0.577 | 0.91489 | 0.1 | 0 | NA | NA |
| Anxiety | C0003467 | GRIA2 | glutamate ionotropic receptor AMPA subunit 2 | 0.573 | 0.692 | 0.99916 | 0.01 | 1 | 2010 | 2010 |
| Anxiety | C0003467 | MATR3 | matrin 3 | 0.631 | 0.538 | 1 | 0.1 | 0 | NA | NA |
| Anxiety | C0003467 | SCN9A | sodium voltage-gated channel alpha subunit 9 | 0.543 | 0.615 | 4.75E-19 | 0.1 | 0 | NA | NA |
| **Anxiety Disorders** | C0003469 | BCHE | butyrylcholinesterase | 0.447 | 0.923 | 1.06E-13 | 0.05 | 5 | 2017 | 2019 |
| Anxiety Disorders | C0003469 | EDIL3 | EGF like repeats and discoidin domains 3 | 0.564 | 0.769 | 0.00014584 | 0.1 | 1 | 2019 | 2019 |
| Anxiety Disorders | C0003469 | FMR1 | FMRP translational regulator 1 | 0.473 | 0.769 | 0.64718 | 0.4 | 16 | 2002 | 2019 |
| Anxiety Disorders | C0003469 | GRIA2 | glutamate ionotropic receptor AMPA subunit 2 | 0.573 | 0.692 | 0.99916 | 0.01 | 1 | 2010 | 2010 |
| Anxiety Disorders | C0003469 | GRIN2B | glutamate ionotropic receptor NMDA subunit 2B | 0.51 | 0.692 | 1 | 0.01 | 1 | 2019 | 2019 |
| **Anxiety symptoms** | C0860603 | BCHE | butyrylcholinesterase | 0.447 | 0.923 | 1.06E-13 | 0.01 | 1 | 2019 | 2019 |
| Anxiety symptoms | C0860603 | FMR1 | FMRP translational regulator 1 | 0.473 | 0.769 | 0.64718 | 0.01 | 1 | 2012 | 2012 |
| **Post-Traumatic Stress Disorder** | C0038436 | BCHE | butyrylcholinesterase | 0.447 | 0.923 | 1.06E-13 | 0.01 | 1 | 2017 | 2017 |
| Post-Traumatic Stress Disorder | C0038436 | FMR1 | FMRP translational regulator 1 | 0.473 | 0.769 | 0.64718 | 0.01 | 1 | 2009 | 2009 |
| Post-Traumatic Stress Disorder | C0038436 | HSP90AA1 | heat shock protein 90 alpha family class A member 1 | 0.411 | 0.923 | 0.86025 | 0.02 | 2 | 2011 | 2018 |
| Post-Traumatic Stress Disorder | C0038436 | PTPN4 | protein tyrosine phosphatase non-receptor type 4 | 0.633 | 0.538 | 0.99663 | 0.01 | 1 | 2018 | 2018 |
| **Stress, Psychological** | C0038443 | FMR1 | FMRP translational regulator 1 | 0.473 | 0.769 | 0.64718 | 0.01 | 1 | 2012 | 2012 |
| Stress, Psychological | C0038443 | GRIN2B | glutamate ionotropic receptor NMDA subunit 2B | 0.51 | 0.692 | 1 | 0.01 | 1 | 2019 | 2019 |

## SUPPLEMENTARY METHODS

### Behavioral data acquisition and analysis

#### Animals

18 adult male Long-Evans rats (260-340 g at the time of surgery, 2-4 months old), were housed in groups of 4 or 5 in large, environmentally-enriched, clear plastic cages (80×60×40cm) before surgery. They were maintained at 21°C in a well-ventilated room with a light/dark cycle of 12h/12h and free access to food and water. Upon arrival in the lab the animals were allowed at least 3 days of acclimatization before being handled daily by the experimenter for at least 5 days prior to surgery. All the experimental procedures were performed in accordance with institutional guidelines and national laws and policies, and were approved by the local Ethics Committee.

#### Surgical implantation of a magnetic base for the Inertial Measurements Unit (IMU)

The IMU was magnetically affixed to the animals’ heads during the experiments. A pair of neodymium disk magnets (S-06-03-N, Supermagnete.com) was glued to the bottom face of the IMU, and another pair was cemented to the skull of the animals with a surgical procedure as described in (Pasquet et al. 2016). Briefly, rats were anesthetized with an intraperitoneal injection of a mixture of ketamine (Imalgene, 180 mg/kg) and xylazine (Rompun, 10 mg/kg). Analgesia was assured by subcutaneous injection of buprenorphine (Buprecare, 0.025 mg/kg). The rats were placed in a stereotaxic frame (Narishige, Japan) and the surgical area was disinfected with povidone-iodine and 70% ethanol. The skull was exposed and gently scraped, and 3% hydrogen peroxide solution was applied. Burr holes were drilled (two in the frontal bone, five in the parietal bones, and one in the occipital bone) and miniature stainless steel screws (Phymep, France) were attached. Self-curing dental adhesive (Super-Bond C&B, Sun Medical) was deposited on the skull. A pair of disk neodymium magnets were glued to a glass/epoxy sheet and were fixed above the screws using self-curing acrylic resin (UNIFAST trad, GC Dental Products Corp.). The skin ridges were sutured in front and at the rear of the implant and the rats were allowed to recover in their home cages for one week. They then were housed individually in standard large cages (58×38×20 cm, DxHxW). After recovery. rats were placed under mild food restriction (~17 g/day, adjusted to allow for a 15 g weight gain per week until reaching 400 g, and then to maintain this weight) to ensure proper motivation for foraging during the sessions in the open field.

#### Behavioral apparati and protocol

A was a transparent plexiglass box (37 × 45 × 37 cm, DxHxW) placed in a noise attenuating cubicle (Med Associates, USA). The floor consisted of 25 stainless steel rods (0.6 cm in diameter) connected to a constant-current scrambled shock generator (ENV-414, Med Associates, USA). C was a custom built PVC cubicle (40 × 40 × 40 cm) of dimensions similar to A, but with gray walls and a solid black floor. The open field could assume two different configurations, B and B’, and waw 250 x 150 cm with 70 cm high walls. While the walls and the floor were composed respectively of PVC and sheets of rubber in both B and B’, the two configurations were different with respect to surface textures, distal cues visible in the room, and the identities of the objects (small models of Parisian and Roman monuments; ~20×20×20cm each) within the open field. In B and B’, to enhance the ethological relevance of the apparatus, we allowed foraging for food pellets (20 mg, MLabRodent Tablet, TestDiet) released from two ceiling-mounted automatic distributors (Camden Instruments, UK) every 20-40 s in each session. The conditioning and fear renewal test chambers were located in differently configured rooms on different floors of the same building. In all environments, behavior was recorded from side-mounted cameras. In A and C, video was captured with a camcorder (Sony Handycam HDR-CX280) and, in B and B’, with 4 video cameras (Basler Aca 2500-60) synchronized with Streampix 6 (Norpix, Canada). In B and B’, video was also recorded at 30 Hz from above the open fields with a webcam (Logitech C920). The positions of red LEDs mounted on the IMU were tracked with Dacqtrack software (Axona, UK). Auditory cues (CS) were 20 s continuous pure tones at 2 kHz (62-68 dB) controlled by a Power1401 interface (CED, UK). This interface also controlled a flashing red LED (invisible to the animals) that helped synchronize position data acquisition. Animals underwent 3 habituation sessions in B and 2 in B’. Half of the animals underwent extinction in B and the others in B’. On day 14, the last day of extinction, half of the animals in B were switched to B’ and vice versa. No difference in any of the behavioral measures was observed among animals who switched configuration on day 14 and those that did not, and their data were pooled.

#### Inertial signals acquisition and processing

The inertial measurement unit (IMU) is a small (19×13×13 mm) and lightweight (6 g) device (Pasquet et al. 2016). The IMU employs Bluetooth wireless communication and the synchronization was assured by an infrared (IR) antenna that captured IR pulses emitted regularly at 0.5 Hz and recorded along with inertial data by the IMU (Pasquet et al. 2016). Sensitivity of the IMU was set to its maximum (±2 g and ±250°/s) and the sampling rate was 300 Hz. Preprocessing of inertial signals was perfomed with custom scripts in R. The head orientations were computed via low-pass filtering of all of the accelerometer signals (Pasquet et al. 2016). A second-order Butterworth low-pass filter with a cut-off frequency of 2 Hz approximated gravity components.

#### Automatic scoring of behavior

All of the behavioral data presented in the figures result from automated scoring pipeline. A supervised deep learning algorithm scored the grooming, rearing, and other foraging/exploratory behaviors. The training and test sets were each derived from 98 minutes of video recordings from two extinction sessions for three randomly selected animals. To create the training and test data sets, an experienced experimenter manually scored behaviors. Grooming was characterized by repetitive motion of the animals’ head, and of the forelimbs to its muzzle and whiskers (face grooming) or body (body grooming). Rearing was characterized as epochs when the animal was standing on its hind limbs. All other time bins were scored as ‘foraging/exploratory’ and mostly included stationary activity, risk assessment, slow locomotion, and foraging behaviors. A three-layer neural network with a convolutional layer and a fully connected hidden layer was implemented with custom scripts and built-in functions of the parallel computing toolbox in Matlab (Mathworks, USA). The behavioral measures provided to the network included: videodetected head position, accelerometer and gyroscope raw signals smoothed using a zero-phase lowpass filter, and the movement frequency power spectrum obtained by wavelet analysis of each gyroscope channel using the WaveletComp package in R that yielded a spectrogram of 3 bands, each 3 Hz wide, spanning from 0.1 to 9 Hz. All signals were downsampled to 30 Hz and binned in 2 s windows with 200 ms overlap. Note that time bins previously classified as darting, freezing or object exploration (as defined above) were excluded from this analyses: the network thus only scored grooming, rearing vs. other behaviors. The network was trained on the manually scored behavior, and the robustness and accuracy of the classification was assessed on the test set by reiterating the training and test process 1000 times. The repeated iterations of the algorithm diverged by 3.8% on average and errors tended to be concentrated in the same 3.9% of bins (**Supp. Fig. 7b-d**). Given the lack of differences between the different iterations, one iteration was randomly selected to classify the entire dataset.

#### Behavioral data analysis

The clustering and all statistical analyses were performed in Matlab (MathWorks, Natick, MA), using the Freely Moving Animal Toolbox (http://fmatoolbox.sourceforge.net) and custom written programs. Behavioral categories whose weights exceeded Otsu’s threshold (Otsu 1979) were considered as significantly contributing to the corresponding principal component. Otsu’s treshold selection is a nonparametric and unsupervised method to set a treshold to extract relevant values from background activity. In figure 3j, to increase the temporal resolution of behavioral cluster separation analysis we separated CS from post-CS in order to represent each session by 6 different intervals. For descriptive statistics, behavioral data was represented with boxplots where the central bar indicates the median, the bottom and top edges indicate the 25th and 75th percentiles; whiskers extend to the most extreme data points, excluding outliers. Datapoints were considered as outliers if they were greater than *q*3 + 2.7*σ*(*q*3-*q*1) or less than *q*1–2.7*σ*(*q*3–*q*1); where *σ* is the standard deviation, and q1 and q3 are, respectively, the 25th and 75th percentiles of the sample data. To represent overall changes in behavioral variance across the protocol (Fig. 2a), the values of variance for the different behavioral classes for each training stage were pooled and fold change was computed relative to the average value of the previous session.

### Transcriptomic analysis

#### Tissue sampling

About 3 hours after the end of the renewal protocol, rats were anesthetized with isoflurane in an induction box, then euthanized with a pentobarbital overdose (160 mg/kg, i.p.). Brains were immediately removed and frozen at −40°C in isopentane for 35 s and stored at −80°C until sectioning. Brains were sectioned (100μm) and the ventromedial prefrontal cortex (vmPFC) in sections from 4 to 2.7 mm rostral from bregma was micro-punched (0.75 mm punch diameter), bilaterally with a probe centered in the middle of the cingulate cortex areas 32 ventral and 25 (Paxinos and Watson 2013). RNA was extracted using Trizol (Invitrogen) and further purified using the Direct-zolTM RNA MiniPrep (Zymo Research). Genome-wide transcriptional profiling was performed for seven rats in each group.

#### Genome sequencing

Library preparation and Illumina sequencing were performed at the Ecole Normale Supérieure genomic core facility (Paris, France). Messenger (polyA+) RNAs were purified from 0.1μg of total RNA using oligo(dT). Libraries were prepared using the strand specific RNA-Seq library preparation TruSeq Stranded mRNA kit (Illumina). Libraries were multiplexed by 14 on 4 high throughput flowcell lanes. A 75 bp single read sequencing was performed on a NextSeq 500 (Illumina). A mean of 2.298 ×10^7^±3.189 passing Illumina quality filter reads was obtained for each of the 14 samples. The analyses were performed using the Eoulsan pipeline (Jourdren et al. 2012), including read filtering, mapping, alignment filtering, read quantification, normalisation and differential analysis. Before mapping, poly N read tails were trimmed, reads ≤40 bases were removed, and reads with quality mean ≤30 were discarded. Reads were then aligned against the Rattus norvegicus genome from Ensembl version 96 using STAR (version 2.7.2d) (Dobin et al. 2013). Alignments from reads matching more than once on the reference genome were removed using Java version of samtools (Li et al. 2009). To compute gene expression, Rattus norvegicus GTF genome annotation version 96 from Ensembl database was used. All overlapping regions between alignments and referenced exons were counted using HTSeq-count 0.5.3 (Anders et al. 2015). The sample counts were normalized using DESeq2 1.8.1 (Love et al. 2014). Statistical treatments and differential analyses were also performed using DESeq2 1.8.1.

#### Differentially expressed genes analysis

Transcriptomics data analysis was performed using routines written in R. Differentially expressed genes (DEGs) were defined by an absolute value of log2 of fold change > 1, an adjusted p-values after Bejamini & Hochberg correction (Benjamini and Hochberg 1995) of < 0.01, and mean total counts for all the conditions > 10. In order to assure stringent selection, further correction was performed non-parametrically with a shuffling technique. We randomly permuted 3431 times (for all possible combinations of two groups of 7 subjects) the assignment of BH corrected p-values to the two groups. For each gene, we computed the number of times the difference in gene expression between the two groups in shuffled data was equal or greater than the difference observed. This was < 0.02 in all cases, and we therefore retained all DEGs. Volcano plots and heatmaps for visualization were generated using the Enhanced Volcano and ComplexHeatmap R packages (Gu et al. 2016; Blighe and RS 2019). Gene annotation and enrichment analysis on the 238 DEGs was performed using Metascape with a minimum count of 5 and default parameters otherwise (http://metascape.org/) (Zhou et al. 2019). All genes in the genome have been used as the enrichment background. Among the DEGs, 190 genes were detected in the ontology sources GO Biological Processes. P-values were calculated based on the accumulative hypergeometric distribution, and q-values were calculated using the Benjamini-Hochberg procedure to account for multiple testing. Protein-protein interactions analysis between the DEGs was performed using STRING (Search Tool for Recurring Instances of Neighboring Genes, v 11.0 https://string-db.org/) (Szklarczyk et al. 2016). Interaction sources include all sources available in STRING. Disconnected nodes are not displayed in Figures. Minimum required interaction score was set to a high confidence level (0.7). Disease Gene association between the DEGs and the stress and anxiety related disorders (“Post-Traumatic Stress Disorder”,”Anxiety Disorders”, “Anxiety symptoms”, “Anxiety”, “Anxiety and fear”, “Abnormal fear/anxiety-related behavior”, “Stress, Psychological”) was performed using DisGeNET database (Piñero et al. 2020).

## Data and code availability

No restriction on data availability applies to this study. Upon publication, the RNASeq gene expression data and raw fastq files will be available on a GEO repository (www.ncbi.nlm.nih.gov/geo/) while behavioral data and code used to perform data analysis will be available on the CRCNS and/or our institutional websites.

## Notes

### Competing Interest Statement

The authors have declared no competing interest.

